# Enhancing Multiplex Genome Editing by Natural Transformation (MuGENT) via inactivation of ssDNA exonucleases

**DOI:** 10.1101/127308

**Authors:** Triana N. Dalia, Soo Hun Yoon, Elisa Galli, Francois-Xavier Barre, Christopher M. Waters, Ankur B. Dalia

## Abstract

Recently, we described a method for multiplex genome editing by natural transformation (MuGENT). Mutant constructs for MuGENT require large arms of homology (>2000 bp) surrounding each genome edit, which necessitates laborious *in vitro* DNA splicing. In *Vibrio* cholerae, we uncover that this requirement is due to cytoplasmic ssDNA exonucleases, which inhibit natural transformation. In ssDNA exonuclease mutants, one arm of homology can be reduced to as little as 40 bp while still promoting integration of genome edits at rates of ~50% without selection *in cis*. Consequently, editing constructs are generated in a single PCR reaction where one homology arm is oligonucleotide encoded. To further enhance editing efficiencies, we also developed a strain for transient inactivation of the mismatch repair system. As a proof-of-concept, we used these advances to rapidly mutate 10 high-affinity binding sites for the nucleoid occlusion protein SlmA and generated a duodecuple mutant of 12 diguanylate cyclases in *V. cholerae*. Whole genome sequencing revealed little to no off-target mutations in these strains. Finally, we show that ssDNA exonucleases inhibit natural transformation in *Acinetobacter baylyi*. Thus, rational removal of ssDNA exonucleases may be broadly applicable for enhancing the efficacy and ease of MuGENT in diverse naturally transformable species.

## INTRODUCTION

Natural transformation is a conserved mechanism of horizontal gene transfer in diverse microbial species. In addition to promoting the exchange of DNA in nature, this process is exploited to make mutant strains in a lab setting. Also, we have recently described MuGENT, a method for multiplex mutagenesis in naturally transformable organisms (1). This method operates under the principle that a subpopulation of cells under competence inducing conditions displays high rates of natural transformation. During MuGENT, cells are incubated with two distinct types of products; one product is a selected marker (i.e. containing an Ab^R^ cassette), which is used to isolate the transformable subpopulation, while the other product is an unselected marker to introduce a mutation (genome edit) of interest without requiring selection *in cis* (i.e. without requiring an antibiotic resistance marker at the edited locus). Upon selection of the selected marker, one can screen for cotransformation of the unselected marker. Under optimal conditions, cotransformation frequencies can be up to 50%. Furthermore, multiple unselected markers can be incubated with cells simultaneously during this process for multiplex mutagenesis.

One requirement for unselected markers during MuGENT is long arms of homology (>2kb) surrounding each genome edit, which are required for high rates of cotransformation. As a result, generating mutant constructs requires laborious *in vitro* splicing of PCR products for each genome edit, which is a major bottleneck for targeting multiple loci during MuGENT. DNA integration during natural transformation is carried out by RecA-mediated homologous recombination. Initiation of RecA-mediated recombination, however, does not require such long regions of homology (2), and theoretically, one long arm of homology should be sufficient to initiate recombination. Therefore, we hypothesized that other factors might necessitate the long arms of homology required for unselected products during MuGENT.

Transforming DNA (tDNA) enters the cytoplasm of naturally transformable species as ssDNA (3). Here, we identify that ssDNA exonucleases inhibit natural transformation in *Vibrio cholerae* and *Acinetobacter baylyi*, two naturally competent species. We exploit this observation to improve MuGENT and perform two proof-of-concept experiments to demonstrate the utility of this method for dissecting complex biological systems and address questions that are impractical using a classical genetic approach.

## MATERIALS AND METHODS

### Bacterial strains and culture conditions

All strains used throughout this study are derived from *V. cholerae* E7946 (4) or *A. baylyi* ADP1 (5). The N16961 strain of El Tor *V. cholerae* was not used in this study because it has a mutation in HapR, which inhibits its natural competence and transformation (6,7). *V. cholerae* strains were routinely grown in LB broth and on LB agar plates supplemented with 50 µg/mL kanamycin, 200 µg/mL spectinomycin, 10 µg/mL trimethoprim, 100 µg/mL carbenicillin, and 100 µg/mL streptomycin as appropriate. *A. baylyi* was routinely grown in LB broth and on LB agar plates supplemented with 50 µg/mL kanamycin or 50 µg/mL spectinomycin as appropriate. A detailed list of all strains used throughout this study can be found in **Table S2**.

### Generation of mutant strains and constructs

Mutant strains were generated by splicing-by-overlap extension (SOE) PCR and natural transformation / cotransformation / MuGENT exactly as previously described (1,8). Briefly, for SOE PCR, primers were engineered to contain overlapping regions in the DNA segments that would be stitched together. All DNA segments were amplified using the high-fidelity polymerase Phusion. Each DNA segment was then gel extracted (to remove template, primers, and any non-specific amplified products). These gel extracted DNA segments then served as template for the SOE PCR reaction using primers that would amplify the final spliced product. For a schematic of SOE PCR see **Fig. S3**. All primers used for making mutant constructs can be found in **Table S3**.

### V. cholerae *transformation assays*

Cells were induced to competence by incubation on chitin (Figure 1 and 2) or via ectopic expression of *tfoX* (P_*tac*_-*tfox*, Figure 3-5) exactly as previously described (1,8). Briefly, competent cells were incubated with tDNA statically at 30°C for ∼5 hours. The tDNA used to test transformation efficiencies throughout this study was ∼500 ng of a linear PCR product that replaced the frame-shifted transposase, VC1807, with an antibiotic resistance cassette (i.e. ΔVC1807::Ab^R^). After incubation with tDNA, reactions were outgrown by adding LB and shaking at 37°C for 2 hours. Reactions were then plated for quantitative culture onto selective media (transformants) and onto nonselective media (total viable counts) to determine the transformation efficiency (defined as transformants / total viable counts).

**Fig. 1.**
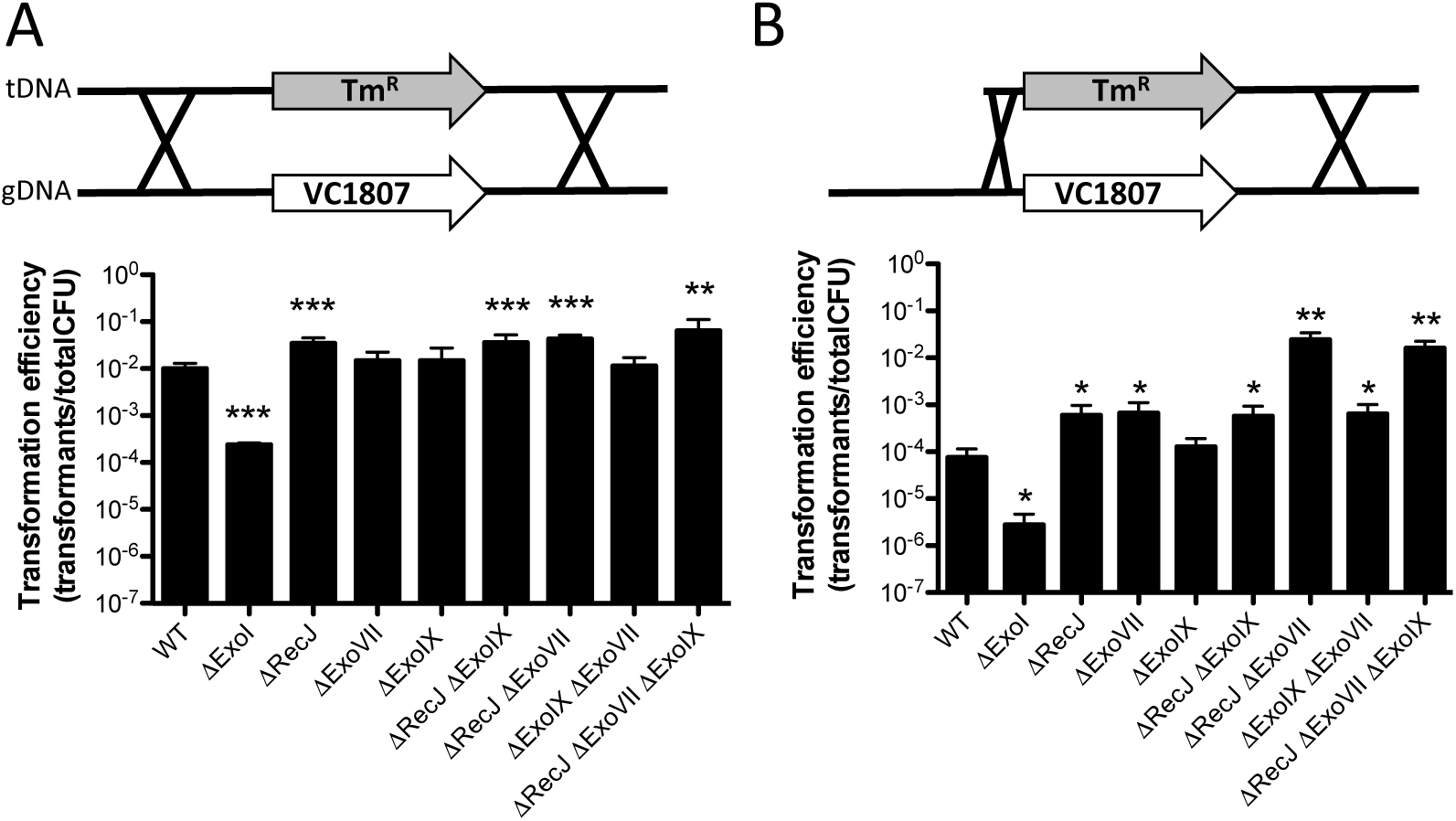
The ssDNA exonucleases RecJ and ExoVII limit natural transformation in V. cholerae. Natural transformation assays of the indicated *V. cholerae* strains with a PCR product as tDNA that has (**A**) 3kb arms of homology on each side of an antibiotic resistance marker (i.e. 3/3 kb) or (**B**) a product where one arm of homology is reduced to just 80bp (i.e. 0.08/3 kb). Data are from at least three independent biological replicates and shown as the mean±SD. Statistical comparisons were made by Student's t-test to the wildtype (WT) strain. *=*p*<0.05, **=*p*<0.01, and ***=*p*<0.001.

**Fig. 2.**
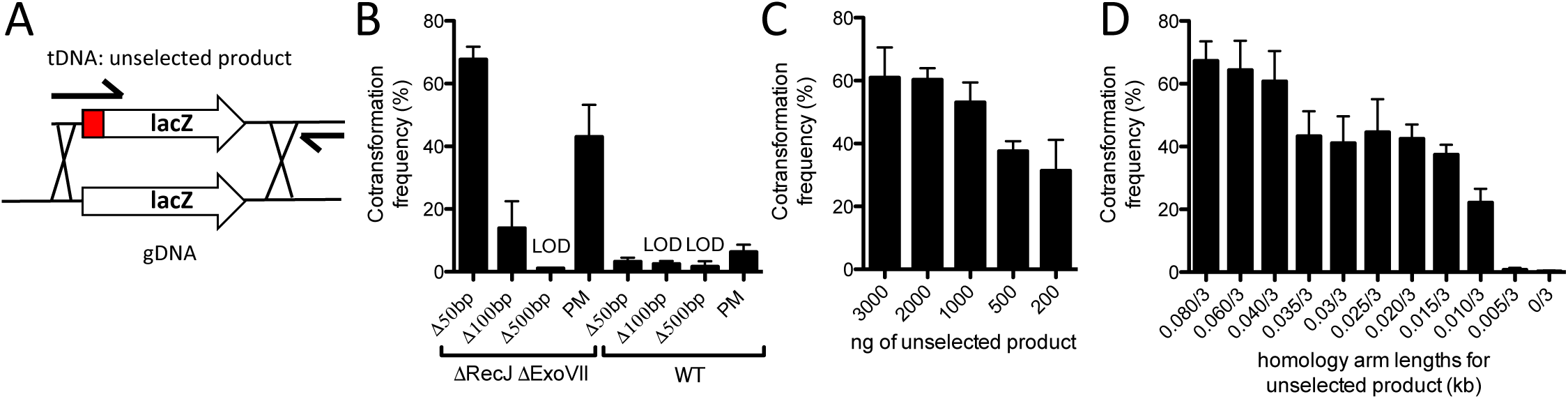
Efficient cotransformation of ssDNA exonuclease mutants using mutant constructs made in a single PCR reaction. (**A**) Schematic indicating how 0.08/3 kb unselected products are generated in a single PCR reaction. The mutation in the unselected product is indicated as a red box and the relative positions of the oligonucleotides used for amplification are highlighted by black arrows. (**B**) Cotransformation assays using 50 ng of a selected product and 3000 ng of an unselected product (all are 0.08/3 kb) into the indicated strain backgrounds. The different unselected products tested generate the indicated type of mutation in the *lacZ* gene. (**C**) Cotransformation assays in a Δ*recJ* Δ*exoVII* mutant using 50 ng of a selected product and the indicated amount of an unselected product (0.08/3 kb) that introduces a 50 bp deletion into the *lacZ* gene. (**D**) Cotransformation assays in a Δ*recJ* Δ*exoVII* mutant using 50 ng of a selected product and 3000 ng of an unselected product (X/3 kb) that introduces a 50 bp deletion into the *lacZ* gene. All data are from at least three independent biological replicates and shown as the mean±SD. LOD=limit of detection and PM=point mutation.

**Fig. 3.**
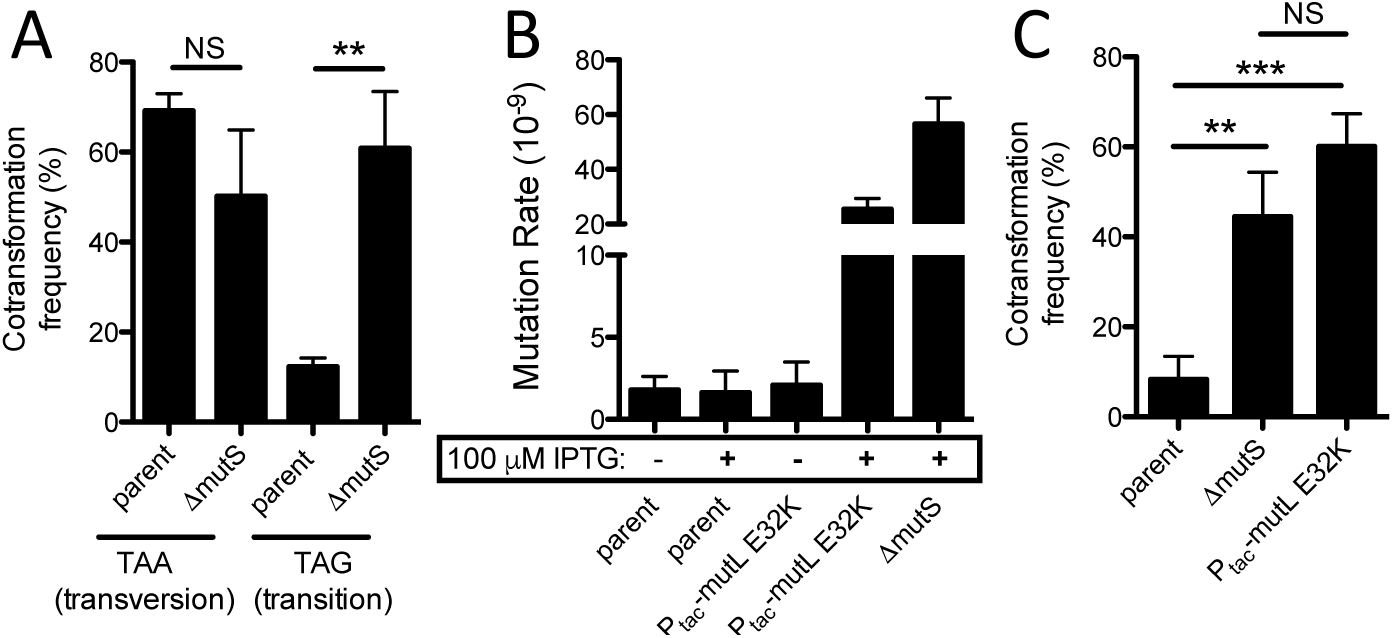
Cotransformation of single point mutations into ssDNA exonuclease mutants is inhibited by MMR and can be overcome by transient expression of a dominant negative allele of MutL. (**A**) Cotransformation assay using an unselected product (0.04/3 kb) to introduce a transversion or transition nonsense point mutation into the *lacZ* gene of the P*_tac_*−*tfoX* Δ*recJ* Δ*exoVII* parent or an isogenic Δ*mutS* mutant. (**B**) Fluctuation analysis for spontaneous resistance to rifampicin to determine the mutation rate of the indicated strains. (**C**) Cotransformation assays in the indicated strains using an unselected product to introduce a transition nonsense point mutation into the *lacZ* gene. All data from **A** and **C** are the result of at least 3 independent biological replicates and data from **B** are from at least 10 independent biological replicates. All data are shown as the mean±SD. **=*p*<0.01, ***=*p*<0.001, and NS=not significant.

**Fig. 4.**
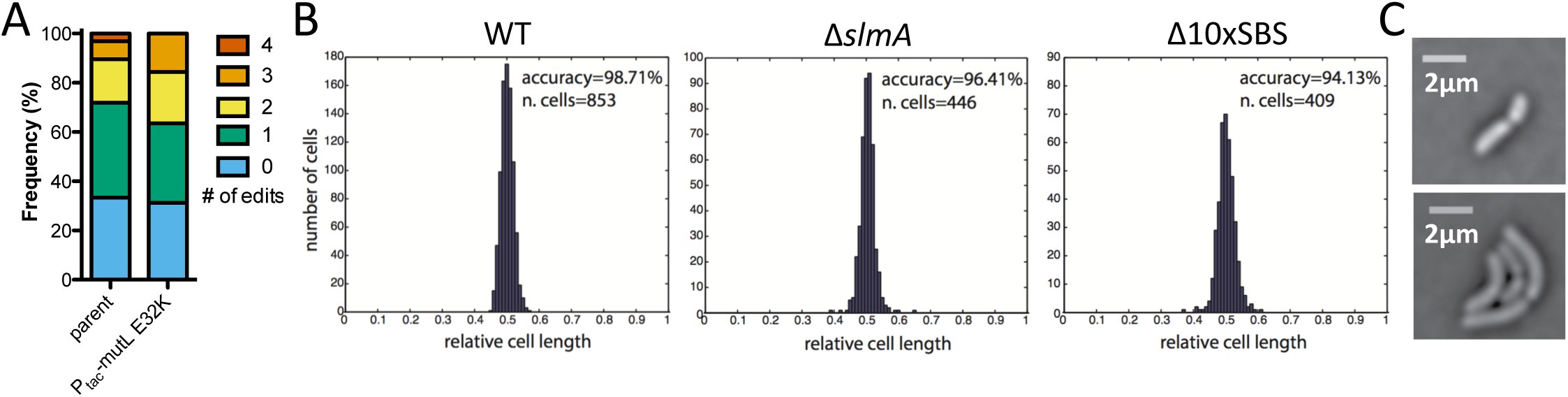
Exo-MuGENT used to rapidly assess the role of DNA binding sites for the nucleoid occlusion protein SlmA. (**A**) Exo-MuGENT was performed in the indicated strains using 100 ng of a selected product and 3000 ng each of 5 distinct unselected products (0.04/3 kb) that introduce point mutations into high-affinity SBSs. Results are shown as the frequency of strains with the indicated number of genome edits following one cycle of MuGENT. (**B**) Cell division accuracy histograms of the indicated strains. Inset values indicate division accuracy, which is defined as the total percentage of cells with a constriction site distant from mid-cell by less than 5%, and the total number of cells analyzed. (**C**) Phase contrast images of rare asymmetric cell division events observed in the Δ10xSBS strain background. Scale bar=2 µm.

**Fig. 5.**
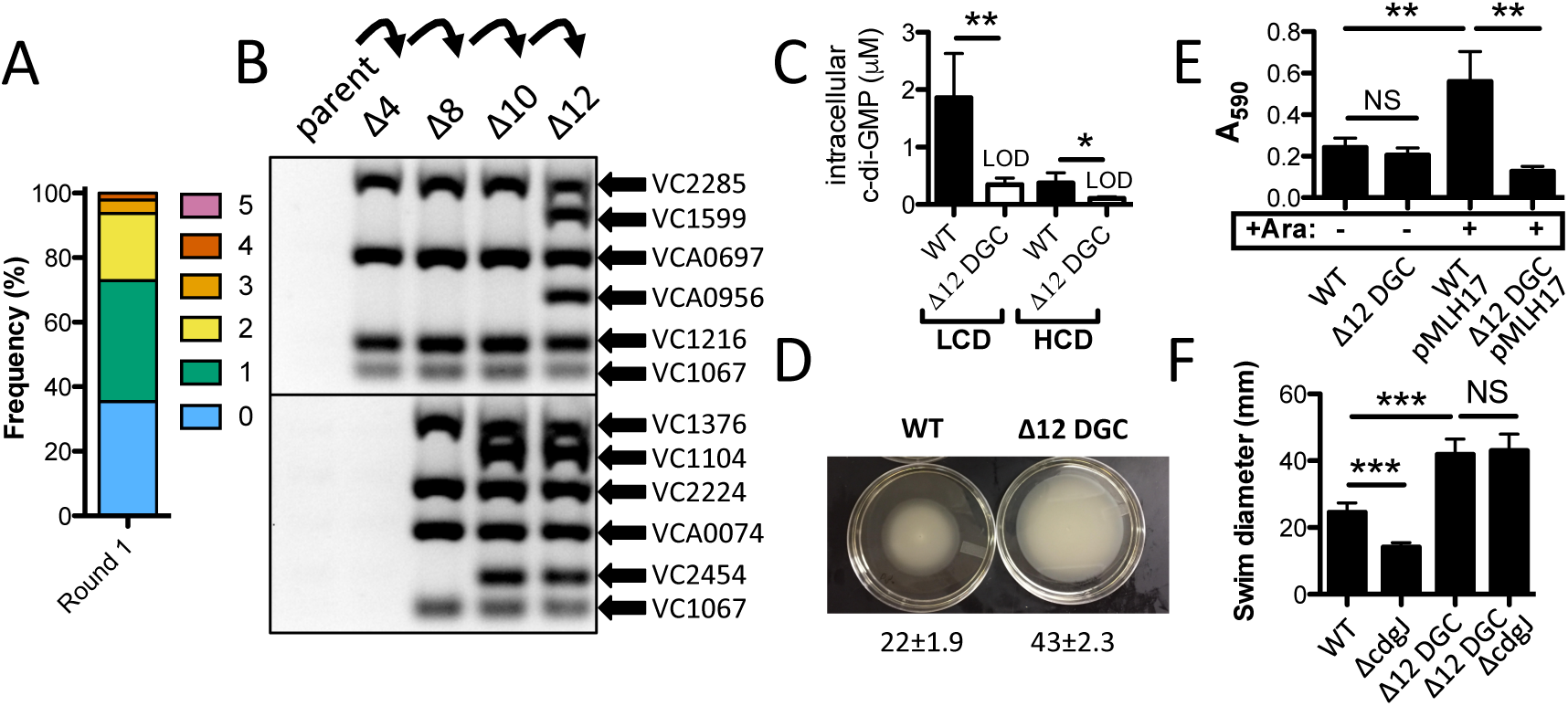
Exo-MuGENT for rapid genetic dissection of diguanylate cyclases. (**A**) Distribution of genome edits in a population of cells following one cycle of Exo-MuGENT using single PCR mutant constructs (0.04/3 kb) that target 12 DGC genes for inactivation. (**B**) MASC-PCR gel for DGC genome edits in intermediate strains leading to generation of the Δ12 DGC mutant. (**C**) Intracellular c-di-GMP concentration of the indicated strains at low cell density (LCD) and high cell density (HCD). Data are from five independent biological replicates and shown as the mean±SD. (**D**) Representative image of a swim assay of the indicated strains with quantification of the swim diameter from six biological replicates shown below as the mean±SD. (**E**) Biofilm assay for the indicated strains. pMLH17 harbors an arabinose-inducible copy of the *vpsR* gene. (**F**) Swim assays of the indicated strains. Data are from at least 8 independent biological replicates and shown as the mean±SD. *=*p*<0.05, **=*p*<0.01, ***=*p*<0.001, and NS=not significant.

### *MuGENT/Cotransformation in* V. cholerae

Cells were induced to competence exactly as described above for transformation assays. Competent cells were incubated with ∼50 ng of a selected product (generally ΔVC1807::Ab^R^) and ∼3000 ng of each unselected product unless otherwise specified. Cells were incubated with tDNA and plated exactly as described above for transformation assays.

Mutations were detected in output transformants by MASC-PCR, which was carried out exactly as previously described (1,9). See **Table S3** for a list of all primers used for MASC-PCR. To generate the Δ10xSBS and Δ12 DGC mutants, the most highly edited strain obtained in the first cycle of MuGENT (from 48 screened) was subjected to an additional round of MuGENT using mutant constructs distinct from those integrated in the first cycle. This process was iteratively performed until all genome edits were incorporated (between 3 and 4 cycles were required for all mutants generated in this study).

To repair the P*_tac_*−*tfoX, recJ, exoVII*, P_*tac*−mutL_ E32K, and *lacZ*::*lacIq* mutations in the Δ10xSBS and Δ12 DGC mutants, strains were subjected to MuGENT to revert these mutations using mutant constructs amplified from the wildtype strain. These unselected products contained 3/3 kb arms of homology. Reversion of these mutations was confirmed by MASC-PCR and through whole genome sequencing.

### Fluctuation analysis for determining mutation rates

Fluctuation analysis for each strain tested was performed by inoculating 10^3^ cells into 10 parallel LB cultures and growing overnight at 30°C for exactly 24 hours. Then, each reaction was plated for quantitative culture on media containing 100 µg / mL rifampicin (to select for spontaneous rifampicin resistant mutants) and onto nonselective media (to determine the total viable counts in each culture). Mutation rates were then estimated using the Ma-Sandri-Sarkar Maximum Likelihood Estimator (MSS-MLE) method using the online FALCOR web interface (10,11).

### Whole genome sequencing

Sequencing libraries for genomic DNA for single end 50 bp reads were prepared for sequencing on the Illumina HiSeq platform exactly as previously described (12). Data were then analyzed for single nucleotide variants and small (<5bp) indels relative to a reference genome using the CLC Genomics Workbench (Qiagen) exactly as previously described (13). For paired end sequencing, genomic DNA libraries were prepped using an NEBNext Ultra kit according to manufacturer's instructions and sequenced on the Illumina MiSeq platform (2 × 300 bp reads). Data were analyzed for structural variants (inversions / deletions), large deletions, and point mutations using CLC Genomics Workbench (Qiagen).

### RNA-seq analysis

RNA was purified using Trizol reagent (ThermoFisher) and then sequencing libraries were prepared using RNAtag-Seq exactly as previously described (14). Reads obtained were mapped to the N16961 reference genome (accession: NC_002505 and NC_002506) and analyzed using the Tufts University Galaxy server (15). Normalized transcript abundance was measured by aggregating reads within a gene and then normalizing for the size of the gene.

### Image analysis for division accuracy and FtsZ cell cycle choreography

Cells were grown in M9 minimal medium supplemented with 0.2% fructose and 1 µg/mL thiamine to exponential phase and then spread on a 1% (w/v) agarose pad of the same medium for microscopic analysis. Septum accuracy was determined in snapshot cell images acquired using a DM6000-B (Leica) microscope and analyzed using MicrobeTracker. *ftsZ-RFPT* was introduced at the *lacZ* locus and expressed from the arabinose promoter using 0.02% of L-Arabinose. FtsZ-RFPT. FtsZ-RFPT cell cycle choreography was determined in time-lapse experiments, the slides were incubated at 30 °C and images acquired using an Evolve 512 EMCCD camera (Roper Scientific) attached to an Axio Observe spinning disk (Zeiss). Image analysis was done as previously described (16,17), in the cell cycle representations colors were assigned to the maximal and minimal fluorescence intensity projections observed at each time point of the time-lapse.

### Assaying intracellular c-di-GMP

To determine intracellular c-di-GMP, overnight cultures were back diluted (1:1000) into 2 mL cultures and grown to the appropriate OD_600_ (LCD: 0.200 – 0.300, HCD: ∼1.00). An aliquot (1.5 mL) of each culture was centrifuged at 15,000 rpm for 30 s and decanted. The remaining pellet was resuspended in 100 µL of cold extraction buffer (40% acetonitrile–40%methanol–0.1Nformic acid) and incubated at −20˚C for 30 min. The sample was centrifuged at 15,000 rpm for 10 min and the supernatant was collected. The solvent was evaporated via a vacuum manifold and the pellet was stored at −80˚C. The pellet was resuspended in 100 µL of HPLC-grade water before quantification of c-di-GMP by an Acquity Ultra Performance liquid chromatography system coupled with a Quattro Premier XE mass spectrometer as previously described (Massie 2012). An 8-point standard curve ranging from 1.9 nM to 250 nM of chemically synthesized c-di-GMP (Biolog) was used to determine [c-di-GMP]. Intracellular concentration of c-di-GMP was estimated as described (18). Because the total c-di-GMP that is measured was divided by a larger intracellular volume at HCD due to higher numbers of cells, this results in HCD having a lower limit of detection for estimated intracellular c-di-GMP than LCD.

### Swim assays

Swimming motility was assayed essentially as previously described (19). Briefly, indicated strains were stabbed into the center of an LB+0.3% soft agar motility plate and then incubated at 30°C for 16 hours. Data is shown as the diameter of the swim radius in mm.

### Biofilm assays

The MBEC Assay System (Innovotech) was used to quantify biofilm formation. An aliquot of each overnight culture (1.5 mL) was pelleted by centrifugation at 15,000 rpm for 3 min. The pellet was washed 3 times with DPBS (1 mL) and resuspended in 1 mL of LB liquid media. The resulting culture was back-diluted to an estimated OD_600_ = 0.001 and aliquoted onto a 96-well MBEC Assay plate (160µL/well, n=5). Strains harboring pMLH17 were induced with 0.2% arabinose and selected with ampicillin (100 µg/mL) before being aliquoted. The plate was incubated at 37˚C for 24 hours shaking at 150 rpm in a humidity chamber to grow biofilms. Biofilm formation was determined by crystal violet staining as previously described (20).

### A. baylyi *transformation assays*

To test transformation efficiency, *A. baylyi* strains were first grown overnight (16-24 hours) in LB medium. Overnight cultures were then spun and resuspended in fresh LB medium to an OD_600_ = 2.0. Then, for each transformation reaction, 50 µL of this culture was diluted into 450 µL of fresh LB medium. Transforming DNA was then added and reactions were incubated at 30°C shaking for 5 hours. The tDNA used to test transformation efficiency in this study replaces a frame-shifted transposase gene, ACIAD1551, with an Ab^R^. Following incubation with tDNA, reactions were plated for quantitative culture onto selective and nonselective media to determine the transformation efficiency as described above.

## RESULTS

### The ssDNA exonucleases RecJ and ExoVII limit natural transformation in V. cholerae

It has previously been shown that high efficiency natural transformation in *V. cholerae* requires long arms of homology surrounding the mutation (1,21). To test this here, we compared rates of transformation using tDNA products containing 3kb arms of homology on each side of an antibiotic resistance marker (i.e. 3/3 kb) or a product where one arm of homology is reduced to just 80bp (i.e. 0.08/3 kb). Consistent with previous studies, we find that the transformation efficiency of the 0.08/3 kb product is ∼100-fold lower than with the 3/3 kb product (Fig. 1). Because RecA should be able to initiate recombination of tDNA with a single long arm of homology, we hypothesized that the reduced rates of transformation observed for the 0.08/3 kb product may be due to DNA endo-or exonucleases that degrade tDNA subsequent to uptake.

One factor previously implicated in limiting natural transformation in *V. cholerae* is the extracellular / periplasmic endonuclease Dns (6). We tested the transformation efficiency of a *dns* mutant using the 3/3 kb and 0.08/3 kb products. While there was a slight increase in the transformation efficiency in the *dns* mutant using the 0.08/3 kb product, this was still at least ∼100-fold lower than when using the 3/3 kb product as tDNA (**Fig. S1**). Thus, Dns is not the main factor that accounts for the relatively poor transformation efficiency of the 0.08/3 kb product. During natural transformation, tDNA translocated into the cytoplasm is single-stranded. Thus, we hypothesized that cytoplasmic ssDNA exonucleases may degrade tDNA after uptake. As a result, rates of transformation for the 0.08/3 kb product would be reduced compared to a 3/3 kb marker since the former has less DNA on one homology arm to serve as a “buffer” for exonuclease degradation. This is consistent with previous work, which demonstrated that extending arms of homology with nonspecific sequences could increase rates of natural transformation (possibly by serving as a buffer for degradation) (21). To test this hypothesis further and identify the ssDNA exonucleases that may be responsible, we targeted the ssDNA exonucleases RecJ, ExoVII, ExoIX, and/or ExoI for inactivation. We found that inactivation of *recJ* and *exoVII* independently resulted in significantly increased rates of natural transformation with the 0.08/3 kb product (Fig. 1). Furthermore, in a *recJ exoVII* double mutant, the transformation efficiency for the 0.08/3 kb product was increased ∼100-fold, which was similar to the efficiency of the wildtype with the 3/3 kb product (Fig. 1). Thus, RecJ and ExoVII both inhibit natural transformation likely by degrading cytoplasmic ssDNA following tDNA uptake. Consistent with RecA-mediated recombination, enhanced transformation efficiency in the *recJ exoVII* double mutant required homology on both sides of the mutation and at least one long arm of homology (**Fig. S2**). It is likely that one long arm of homology is required to initiate homologous recombination at high efficiency during natural transformation.

### High efficiency cotransformation of ssDNA exonuclease mutants with single PCR mutant constructs

Above, we found that as long as there was one long arm of homology (3kb), the other arm of homology could be 80 bp on a selected marker to support high efficiency integration of tDNA. For a selected product (i.e. a product containing a selectable Ab^R^ marker), this has limited utility since generating these mutant constructs still requires *in vitro* DNA splicing of the Ab^R^ cassette to one arm of homology. Exploiting ssDNA exonuclease mutants for mutagenesis would be much more useful if similar arms of homology could be used on unselected products during MuGENT. Normally, to make 3/3 kb mutant constructs for defined deletions or point mutations, the mutation is engineered onto the oligonucleotides used to amplify the upstream and downstream region of homology (**Fig. S3**). Then, *in vitro* splicing of these products generates the final mutant construct. If 80 bp of homology is sufficient in ssDNA exonuclease mutant backgrounds, however, this can be engineered onto the same oligonucleotide used to generate the mutation. This would allow for generation of mutant constructs with 0.08/3 kb homology in a single PCR reaction (Fig. 2A and **Fig S3**).

To determine if mutant constructs generated in a single PCR reaction could be used for MuGENT in ssDNA exonuclease mutant backgrounds, we first tested integration of one unselected product (also called cotransformation). The unselected genome edits tested introduced a 50 bp deletion, 100 bp deletion, 500 bp deletion, or a transversion nonsense point mutation into the *lacZ* coding sequence, which provided a simple readout for cotransformation on X-gal containing medium. All point mutant and deletion unselected products were generated in a single PCR reaction by engineering the mutation onto the same oligonucleotide encoding the 80 bp arm of homology (**Fig. S3**). Single unselected products were incubated with a competent population along with a selected marker that replaces the frame-shifted transposase VC1807 with an Ab^R^ marker. We found that cotransformation rates for all of the unselected products tested were either at or near the limit of detection in the wildtype strain background (**Fig. 2B**). In the *recJ exoVII* mutant, however, a 50bp deletion and a transversion point mutation could be integrated at cotransformation rates of ∼50% (**Fig 2B**). Single PCR mutant constructs could not promote high efficiency cotransformation of 100 bp or 500 bp deletions though (**Fig. 2B**). The concentration of unselected product required for high rates of cotransformation are ∼1000ng, which is ∼3 times lower than that required in the original MuGENT protocol (Fig. 2C) (1). This is likely due to reduced degradation of cytoplasmic tDNA in the ssDNA exonuclease mutant background. We also sought to determine the shortest length of homology required to facilitate efficient cotransformation. To that end, we tested unselected products with reduced lengths of the “short” oligonucleotide encoded arm of homology. We found that a short arm of homology of even ∼10-15 bp allowed for efficient cotransformation, however, the highest rates observed required ∼40 bp of homology (Fig. 2D). Thus, cotransformation in ssDNA exonuclease mutant backgrounds allows for high efficiency integration of unselected products generated via a single PCR reaction without any *in vitro* DNA splicing.

### Subverting MMR in ssDNA exonuclease mutant backgrounds using a dominant negative allele of MutL

Single-stranded exonucleases, including RecJ and ExoVII, participate in methyl-directed mismatch repair (MMR) by excising and degrading the mutated strand (22). Thus, while a *recJ exoVII* double mutant may allow for highly efficient natural transformation, an off-target effect may be an increased rate of spontaneous mutations via reduced MMR activity. To test this, we performed fluctuation tests for spontaneous resistance to rifampicin to determine the mutation rates of select ssDNA exonuclease mutants. We found that the *recJ exoVII* double mutant had a mutation rate similar to the wildtype (**Fig. S4**), indicating intact MMR activity in this mutant background.

Conversely, MMR can inhibit natural transformation in many bacterial species (3). We therefore tested if MMR inhibited natural transformation of point mutations in ssDNA exonuclease mutants using 0.04/3 kb mutant constructs. To induce transformation in these experiments, strains contained the master regulator of competence, *tfoX*, under the control of an IPTG-inducible P_*tac*_ promoter (8). Generally, transition mutations are efficiently repaired by the MMR system, while transversion mutations are poorly recognized (23,24). Cotransformation of a transition point mutation into the parent *recJ exoVII* mutant is significantly reduced compared to a transversion point mutation (Fig. 3A). Conversely, both types of point mutations are integrated at equal rates in a Δ*mutS* MMR-deficient background (Fig. 3A). Thus, these results indicate that MMR can inhibit the integration of point mutations in ssDNA exonuclease mutant backgrounds when using 0.04/3 kb mutant constructs.

While inactivation of MMR allowed for highly efficient integration of genome edits with point mutations, performing mutagenesis in MMR deficient backgrounds is not optimal, as these strains would accumulate a large number of off-target mutations. So next, we sought to generate a strain where we could transiently inactivate the MMR system. Recently, a dominant-negative allele of MutL (E32K) was used to transiently inactivate MMR during multiplex automated genome engineering (MAGE) in *Escherichia coli* (25). We generated a strain of *V. cholerae* where expression of *mutL* E32K was driven by an IPTG-inducible P_*tac*_ promoter in the Δ*recJ* Δ*exoVII* P_*tac*_−*tfoX* mutant strain background. As expected, the spontaneous mutation rate of this strain was similar to the parent strain in the absence of IPTG, while it approached the mutation rate of an MMR-deficient mutant in the presence of 100 µM IPTG (Fig. 3B), indicating that overexpression of *mutL* E32K, indeed, functionally inactivates the MMR system. Next, we tested the cotransformation efficiency of a genome edit with a transition point mutation into this transient mutator strain. In the P_*tac*_-*mutL* E32K background, we observed remarkably high cotransformation efficiencies for a transition point mutation, which was equivalent to the cotransformation efficiency observed in an MMR deficient background (Fig. 3C). Cells only undergo 1-2 generations of growth in the presence of IPTG in these experiments (i.e. conditions where MMR is transiently inactivated); thus, the number of spontaneous mutations that will be introduced during this procedure should be minimal, which was tested further below.

### *Exo-MuGENT for dissection of SlmA function in* V. cholerae

Having successfully introduced one unselected product into ssDNA exonuclease mutants using a 0.04/3 kb mutant construct, we next wanted to test MuGENT with multiple 0.04/3 kb unselected products in ssDNA exonuclease mutant backgrounds. We refer to this new approach as Exo-MuGENT. In a first proof-of-concept experiment, we used Exo-MuGENT to introduce multiple unselected products containing specific point mutations into the genome. As a target, we studied the nucleoid occlusion protein SlmA, which binds to specific sequences in the genome called SlmA binding sites (SBSs). DNA-binding by SlmA allows it to inhibit FtsZ polymerization over the nucleoid, which is required for proper placement of the division site in *V. cholerae* (17). Indeed, inactivation of *slmA* results in mislocalization of FtsZ during the cell cycle (17). There are 79 SBSs in the *V. cholerae* chromosome (17) with varying affinities for SlmA. We decided to initially determine the role of the highest-affinity SBSs on nucleoid occlusion activity. To test this, we performed Exo-MuGENT to mutate conserved residues in the 10 SBSs with the highest affinity for SlmA (17). The mutant constructs contained homology lengths of 0.04/3 kb, where the 40bp of homology was appended onto the same oligonucleotide used to introduce the point mutations into the SBSs. Using this approach, we first incubated 5 unselected products with a population of competent cells to target 5 distinct SBSs for mutagenesis. This was performed in both the parent strain background (Δ*recJ* Δ*exoVII* P_*tac*_-*tfoX*) and in the P*tac*− *mutL* E32K transient mutator strain (Δ*recJ* Δ*exoVII* P*tac*-*tfoX* P*tac*−*mutL* E32K). Surprisingly, Exo-MuGENT yielded highly complex mutant populations in both strain backgrounds with a significant fraction (10-15%) of both populations containing 3-4 genome edits (Fig. 4A). Thus, while MMR may limit integration of unselected products with a single point mutation (Fig. 3C), Exo-MuGENT with multiple unselected products that introduce a large number of point mutations may overwhelm the MMR system. This phenomenon is also observed in other naturally competent species (26). Thus, depending on the application, both the parent and transient mutator strain backgrounds can be used to generate highly edited strain backgrounds and complex mutant populations.

Next, we took the strains with the greatest number of genome edits in the first cycle of Exo-MuGENT and then subjected each to additional cycles of editing. As for the original MuGENT protocol, the selected products used in each cycle of Exo-MuGENT swapped the antibiotic resistance marker at the selected locus (i.e. introduction of a new resistance cassette replaced the endogenous one), which facilitates recycling of resistance markers throughout the procedure. We selected the most highly edited strain at each cycle and continued until all 10 genome edits were incorporated. This process took 3 cycles of Exo-MuGENT in the the P_*tac*_−*mutL* E32K strain background and 4 cycles in the parent strain background. We then repaired the *recJ, exoVII*, P_*tac*_−*mutL* E32K, and P_*tac*_−*tfoX* mutations in both strains backgrounds to generate SBS edited strains (referred to as Δ10xSBS) that were isogenic with our wild type strain (see Methods for details). Repair of these alleles was at least as efficient as introduction of the SBS genome edits and took only 1-2 cycles of Exo-MuGENT to accomplish. Thus, these strains were subjected to 14 distinct genome edits by MuGENT in ssDNA exonuclease mutant backgrounds.

One concern when performing multiplex mutagenesis is the accumulation of off-target mutations. The original MuGENT protocol resulted in little to no off-target mutations, even in strains with 13 genome edits (1,27). To determine if this was also true for Exo-MuGENT, we sequenced the whole genomes of the Δ10xSBS strains obtained in the P_*tac*_-*mutL* E32K and parent strain backgrounds (See Methods for details) and analyzed this data for the presence of off-target point mutations and/or small indels relative to the parent strain. For the Δ10xSBS mutant in the parent strain background, we identified two non-synonymous point mutations in the *alaS* and *rdgC* genes, which were not within the mutant constructs of any of the SBSs targeted by our approach. So, these were likely spontaneous point mutations in this strain. In the P_*tac*_-*mutL* E32K strain background, despite transient suppression of mismatch repair during mutagenesis, we found no off-target mutations in the Δ10xSBS mutant. As a result, the latter strain was characterized further for its impact on cell division licensing.

First, we found that FtsZ localization throughout the cell cycle in the Δ10xSBS mutant was largely similar to the wildtype, indicating that these 10 high-affinity SBSs do not account for the majority of SlmA-dependent nucleoid occlusion activity (**Fig. S5**). Cell division accuracy can be assessed by measuring the faithful placement of the septum at midcell in dividing cells. In the Δ10xSBS mutant we did observe a small but significant reduction in cell division accuracy (Fig. 4B and C). Thus, these 10 SBSs may play a subtle role in the proper placement of the division site in *V. cholerae*. Cumulatively, these results demonstrate that Exo-MuGENT (i.e. the use of ssDNA exonuclease mutants for MuGENT with single PCR mutant constructs) dramatically simplifies the procedure and provides an efficient means to rapidly generate highly edited bacterial genomes with little to no off-target mutations.

### *Exo-MuGENT for genetic dissection of c-di-GMP biogenesis in* V. cholerae

Next, we attempted to use Exo-MuGENT to introduce multiple deletions into the genome in a second proof-of-concept experiment. The targets in this second experiment are the genes required for biogenesis of cyclic di-GMP (c-di-GMP). This secondary messenger regulates the transition between motile and sessile lifestyles in microbial species and is generated by the action of diguanylate cyclases (DGCs) (28). Generally, increased levels of c-di-GMP correlate with decreased motility and enhanced biofilm formation, while decreased c-di-GMP levels are correlated with increased motility and decreased biofilm formation. There are 41 distinct DGCs in *V. cholerae* (29). Since c-di-GMP is readily diffusible, this compound can mediate a global regulatory response (low specificity signaling), however, it has been demonstrated in *V. cholerae* that specific DGCs mediate distinct outputs that are independent of the cellular c-di-GMP concentration (18). This argues a high-specificity of signaling via distinct DGCs. Also, with such a large number of DGCs there is the possibility that enzymes within this class are genetically redundant. To begin to genetically dissect this system, we decided to inactivate the 12 DGCs that are most highly expressed (as determined by RNA-seq) or that were previously implicated in c-di-GMP-dependent phenotypes during growth in rich medium (**Table S1**) (30,31). Genes were inactivated using mutant constructs that replaced 48 bp of the 5’ end of each gene targeted with a sequence to introduce a premature stop codon (**Fig. S3**). These mutant constructs had 0.04/3 kb of homology, where the 40 bp of homology was appended onto the oligonucleotide used to introduce the mutation. Thus, each unselected product was generated in a single PCR reaction as described above (Fig. 2A and **Fig. S3**). We performed Exo-MuGENT in the *recJ, exoVII*, P_*tac*_-*tfoX* strain with 5 of these unselected products. In a single cycle, ∼60% of the population had one or more genome edits with a small portion (∼6%) of the population having 3-4 genome edits incorporated in a single step (Fig. 5A). The quadruple mutant generated in the first cycle of this process was subjected to three additional rounds of MuGENT with the most highly edited strain at each step being carried over into the subsequent cycle until all 12 of the DGCs targeted were inactivated (Fig. 5B). Then, we repaired the *recJ, exoVII*, P_*tac*_-*tfoX*, and *lacZ*::*lacIq* mutations in this background to make a strain that was isogenic with the wildtype. Despite the 16 genetic modifications, whole genome sequencing via paired-end sequencing (2 × 300 bp reads) revealed that this strain had no off target mutations (including point mutations, inversions, small indels, or large deletions).

In *V. cholerae*, c-di-GMP is produced at higher levels during growth at low cell density (LCD) compared to high cell density (HCD) (32). We found that c-di-GMP levels were at the limit of detection in cell extracts of the Δ12 DGC mutant at both LCD and HCD as measured by liquid chromatography coupled with tandem mass spectrometry (LC-MS/MS), indicating that the 12 DGCs targeted play a major role in producing this second messenger during growth in rich medium (Fig. 5C). Note that the limit of detection at LCD is different from that at HCD in these assays (see Methods for details). Low levels of c-di-GMP are correlated with enhanced motility and decreased biofilm formation in *V. cholerae* (33). Consistent with this, we find that the Δ12 DGC strain displays significantly increased motility on swim agar (Fig. 5D). The E7946 strain used throughout this study is a smooth variant of *V. cholerae* and naturally produces poor biofilms. Therefore, we did not observe a decrease in biofilm formation in the WT compared with the Δ12 DGC strain. However, one transcriptional activator required for extracellular polysaccharide production and biofilm formation is VpsR (34). To activate transcription, VpsR directly binds to c-di-GMP (35). We found that ectopic expression of VpsR in the wildtype background resulted in a significant increase in biofilm formation, presumably by enhancing the activity of the basal levels of c-di-GMP produced (Fig. 5E). VpsR is not, however, sufficient and c-di-GMP is still required for activation of downstream pathways required for biofilm formation (35,36). Consistent with this, we find that VpsR overexpression in the Δ12 DGC strain does not increase biofilm formation (Fig. 5E). Thus, this suggests that some combination of the 12 DGCs targeted generate the c-di-GMP required for VpsR-dependent biofilm formation in rich medium.

The phenotype of the Δ12 DGC mutant on swim agar was more severe than any single DGC mutant strain, suggesting that the 12 DGCs targeted work in concert to additively or synergistically decrease motility (**Fig. S6**). Indeed, redundancy amongst DGCs has been previously reported (30,31). C-di-GMP is degraded by phosphodiesterases (PDE), thus in PDE mutant strains, one would expect elevated levels of c-di-GMP. It was previously shown that inactivation of the PDE *cdgJ* dramatically reduces swimming motility as a result of elevated c-di-GMP-mediated mannose-sensitive hemagglutinin pilus (MSHA) activity (30,37). Inactivation of 4 DGCs in the *cdgJ* mutant resulted in increased swimming motility (due to reducing c-di-GMP levels), however, the swimming motility observed was still significantly lower than an isogenic strain where the 4 DGCs were inactivated in an otherwise wildtype background (30). This suggested that additional DGCs (other than the 4 targeted) might still be generating c-di-GMP, which results in reduced motility in the *cdgJ* mutant background. Consistent with prior work, the *cdgJ* mutant in a wildtype background resulted in a significantly decreased swim radius in our hands (Fig. 5F). In the Δ12 DGC strain background, however, swimming was unaffected by inactivation of *cdgJ* (Fig. 5F). Cumulatively, these results indicate that these 12 DGCs are primarily responsible for production of c-di-GMP in rich medium to promote both swimming motility and biofilm formation in *V. cholerae*.

### *Natural transformation in* Acinetobacter baylyi *is also inhibited by cytoplasmic ssDNA exonucleases*

Another highly naturally competent Gram-negative organism is *Acinetobacter baylyi*. It was previously shown that the ssDNA exonuclease RecJ limits integration of tDNA by homology-facilitated illegitimate recombination (HFIR), but not by truly homologous recombination during natural transformation (38). Homologous recombination, however, was tested in that study by using mutant constructs containing long regions of homology on both sides of the mutation. Indeed, we also find little impact of the ssDNA exonucleases RecJ and ExoX when using tDNA containing long arms of homology (3/3 kb)(Fig. 6). If we use tDNA that contains one short arm of homology (0.08/3 kb), we find that *recJ exoX* double mutants are significantly more transformable than the parent strain, albeit not to the levels observed when using the 3/3 kb product (Fig. 6). This result suggests that ssDNA exonucleases inhibit natural transformation in *A. baylyi* as observed in *V. cholerae*. Thus, rational removal of ssDNA exonucleases may be a viable approach to enhance the efficiency and ease of MuGENT in diverse naturally competent microbial species.

**Fig. 6.**
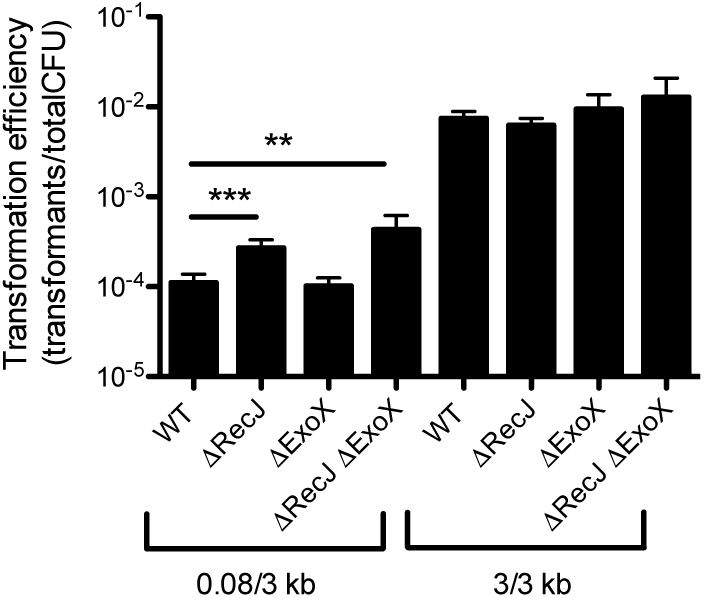
ssDNA exonucleases inhibit natural transformation in Acinetobacter baylyi. (**A**) Transformation assay of the indicated strains using tDNA that contains either 0.08/3 kb arms of homology or 3/3 kb arms of homology as indicated. Data are the result of at least three independent biological replicates and shown as the mean±SD. **=*p*<0.01 and ***=*p*<0.001.

## DISCUSSION

In this study we uncover that cytoplasmic ssDNA exonucleases degrade tDNA to inhibit natural transformation in *V. cholerae* and *A. baylyi*. Through systematic genetic dissection of ssDNA exonucleases in *V. cholerae* we identified that RecJ and ExoVII are likely the main inhibitors of this process. In a *recJ exoVII* double mutant, we found that one arm of homology can be reduced to as little as 10-15 bp and still support high rates of natural transformation. We further exploited this discovery to perform MuGENT in ssDNA exonuclease mutants using mutant constructs that are generated in a single PCR reaction, an advance we have termed Exo-MuGENT. This improvement greatly simplifies the procedure and enhances the scalability of MuGENT in *V. cholerae* to promote generation of highly edited bacterial genomes on even faster timescales. Indeed rational removal of nucleases has also been used to improve recombineering, a distinct genome editing method, in *E. coli* (39). In *A. baylyi* we also found that ssDNA exonucleases inhibit natural transformation. However, rates of transformation with the 0.08/3 kb product in the *recJ exoX* mutant were still lower than when using the 3/3 kb product (Fig. 6). This suggests that other factors (additional ssDNA exonucleases, the MMR system, DNA endonucleases, etc.) may still prevent high efficiency integration of tDNA in *A. baylyi*. Identifying and characterizing these factors will be the focus of future work. One concern during mutagenesis is the introduction of off-target mutations. Indeed, exonucleases are important for MMR activity and *exoVII* mutants have previously been shown to have a hyper-recombination phenotype (40,41). Whole genome sequencing of our most highly edited strain, however, did not identify any off-target mutations. Thus, Exo-MuGENT is a robust method for editing genomes with high efficiency and fidelity.

We performed two proof-of-concept experiments to demonstrate the utility and speed of Exo-MuGENT in *V. cholerae*. First, we mutated 10 high-affinity binding sites for the nucleoid occlusion protein SlmA. This analysis uncovered that these 10 binding sites play a small, yet significant role in accurate cell division. Further mutagenesis of the remaining 69 SBSs may uncover which binding sites are critical for SlmA-dependent nucleoid occlusion. Furthermore, a strain lacking all SBSs would allow for testing which SBSs (or genomic positions) are sufficient to promote nucleoid occlusion, which will be the focus of future work. This experiment also provides a framework for assessing the function of additional DNA binding proteins (e.g. transcription factors, chromosomal macrodomain proteins, partitioning systems, etc.). Using Exo-MuGENT to mutagenize DNA binding sites can rapidly uncover which genetic loci are critical for a particular phenotype. Also, Exo-MuGENT could be used to introduce novel binding sites at disparate genetic loci to determine the role of chromosomal positioning on gene function.

In our second proof-of-concept experiment, we used Exo-MuGENT to inactivate the 12 DGCs that were mostly highly expressed and/or active during growth in rich medium. This resulted in a strain that produced no detectable c-di-GMP. Consistent with this, we find that even when the phosphodiesterase *cdgJ* is inactivated, the Δ12 DGC strain does not display reduced motility, a c-di-GMP dependent phenotype. As a result, we believe this mutant represents a genetic background with the lowest c-di-GMP levels observed in *V. cholerae*. Future work will focus on defining if any of the 12 DGCs studied here are independently sufficient to promote biofilm formation and reduce motility and/or if there are genetic interactions between these genes. Also, we will use Exo-MuGENT to systematically genetically dissect each of the 41 DGCs in *V. cholerae* as well as the 30 genes involved in degradation of this secondary messenger by first making a strain of *V. cholerae* that lacks all of these genes. This analysis could uncover genetic interactions among these loci as well as novel roles for c-di-GMP in *V. cholerae* biology.

Exo-MuGENT provides a significant advance over the original MuGENT protocol. First, mutant constructs for Exo-MuGENT can be made in a single PCR reaction while products for MuGENT require laborious *in vitro* DNA splicing. Thus, mutant constructs for Exo-MuGENT take hours to generate, while those for MuGENT generally take 2 days to prepare (see **Fig. S3** for details). Additionally, failed SOE reactions are not uncommon when making mutant constructs for MuGENT and are burdensome to optimize. The ability to make mutant constructs in a single PCR reaction also allows for this process to be robotically automated; something that is difficult for MuGENT since SOE PCR requires gel extraction of PCR products (**Fig. S3**). This is highly advantagenous for genome scale editing where dozens of genome edits will be generated.

A major drawback to Exo-MuGENT, however, is that a modified strain background must be used (e.g. a P_*tac*_−*tfoX recJ exoVII* mutant), whereas the original MuGENT protocol worked in completely WT *V. cholerae*. In many cases, the *recJ* and *exoVII* mutations may not pose any issue; however, to make strains completely isogenic to wildtype, these mutations must be repaired, which adds 1-2 cycles of Exo-MuGENT to strain construction. Thus, it is most valuable to employ Exo-MuGENT over MuGENT when a large number of genome edits are desired. Each cycle of MuGENT takes ∼3 days to perform, while each cycle of Exo-MuGENT can be carried out in ∼2 days. Thus, for <5-6 genome edits it may be faster to employ MuGENT (assuming mutant constructs can be easily obtained by SOE PCR), while generating 5+ genome edits may warrant the use of Exo-MuGENT. For making highly edited genomes with dozens of mutations, Exo-MuGENT would be highly preferred. For example, making a strain with 50 genome edits by MuGENT would take ∼52 days to complete (assuming that 3 genome edits can be introduced per cycle on average), while at the same editing efficiency this mutant could be made in ∼36 days by Exo-MuGENT.

Exo-MuGENT adds to the continually developing toolbox of methods for synthetic biology in microbial systems. One major advantage of MuGENT over other methods for multiplex genome editing (e.g. MAGE) is the ability to introduce both small and sizable genome edits. The size of genome edits that can be incorporated at high-efficiency by Exo-MuGENT while sizable (∼50 bp), are significantly less than what was possible with the original MuGENT method (∼500-1000 bp genome edits at similar efficiencies). This can still allow for promoter swaps, RBS tuning, gene inactivation, etc. For most applications, the substantial benefit of easily generated mutant constructs (as discussed above) can be a worthwhile tradeoff. MuGENT has now been demonstrated in diverse species including *V. cholerae* (1), *Vibrio natriegens* (42), *Streptococcus pneumoniae* (1), *Helicobacter pylori* (43) and *A. baylyi* (our unpublished results). Thus, Exo-MuGENT may be a viable approach in many of these and possibly other naturally competent organisms.

## ACKNOWLEDGEMENTS

We would like to thank Neil Greene for helpful discussions and assistance with whole genome sequencing. We would also like to thank Tufts TUCF Genomics, the Indiana University CGB, and the Michigan State University Mass Spectrometry Facility for assistance with whole genome sequencing and mass spectrometry, respectively. This work was supported by US National Institutes of Health Grant AI118863 to ABD, GM109259 to CMW, and startup funds from the Indiana University College of Arts and Sciences to ABD. EG and FXB were financially supported by the European Research Council under the European Community's Seventh Framework Programme [FP7/2007-2013 Grant Agreement no. 281590].

## SUPPLEMENTARY FIGURES

**Fig. S1.**
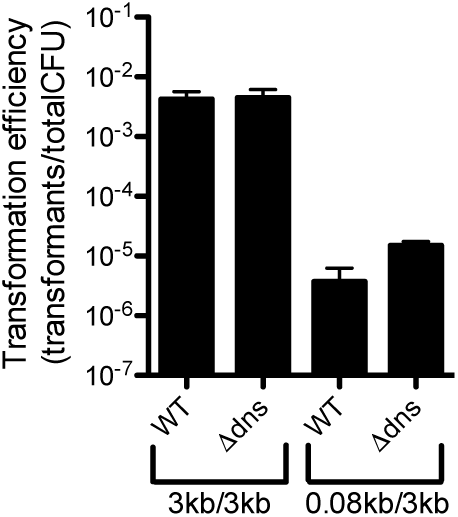
Dns does not inhibit natural transformation of tDNA with one small arm of homology. Natural transformation assay of the indicated strains with tDNA containing the indicated length of homology on either side of the mutation. All data are the result of at least three independent biological replicates and are shown as the mean±SD.

**Fig. S2.**
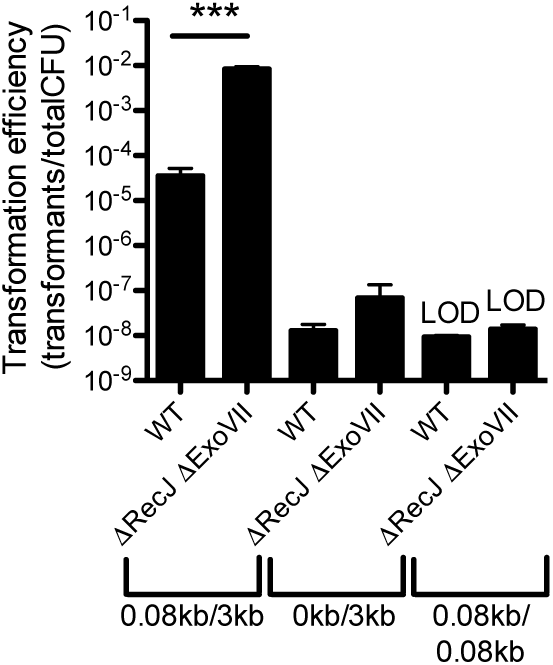
High efficiency transformation requires two arms of homology where at least one arm is long. Natural transformation assay of the indicated strains with tDNA containing the indicated length of homology on either side of the mutation. All data are the result of at least three independent biological replicates and are shown as the mean±SD. ***=*p*<0.001 and LOD=limit of detection.

**Fig. S3.**
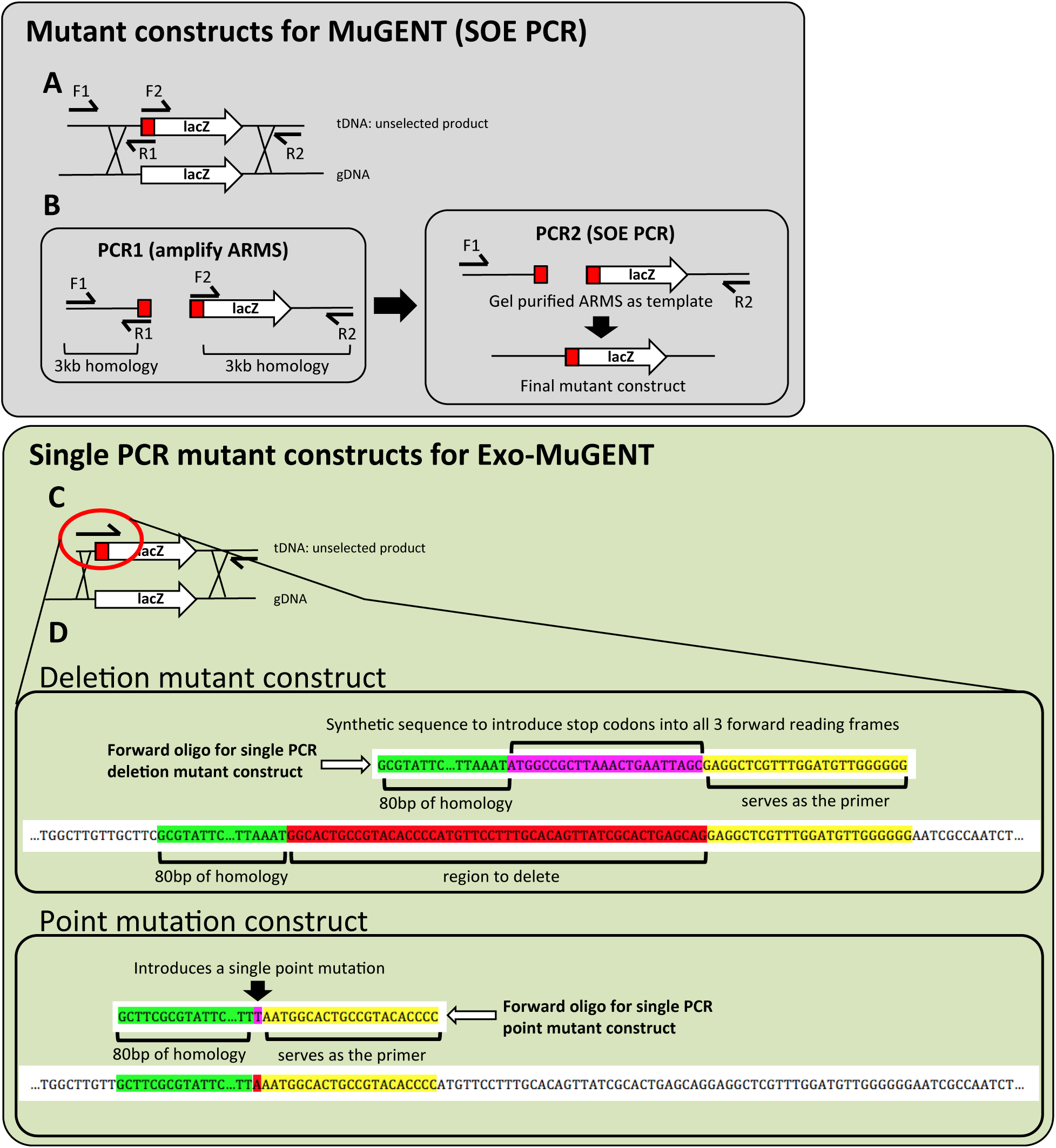
Schematic for generating mutant constructs for MuGENT and Exo-MuGENT. (**A** and **B**) Mutant construct generation for classical MuGENT in WT *V. cholerae*. (**A**) Shows the general overview of how mutant constructs are generated by SOE PCR. (**B**) Schematic for the two distinct PCR steps required for generating mutant constructs. In the first round of PCR, arms of homology are amplified off of genomic DNA with F1/R1 and F2/R2 oligos as indicated. The R1 and F2 oligos are engineered to contain the mutation of interest (deletion, insertion, and/or point mutation), which are highlighted in red in the schematic. The products from the first PCR are then gel purified and serve as template for a second PCR reaction. The overlapping ends of the two ARMS allows them to be spliced together and the final product is amplified with the F1 and R2 primers. (**C** and **D**) Making single PCR mutant constructs for Exo-MuGENT in ssDNA exonuclease mutant strain backgrounds. (**C**) Overview of single PCR mutant constructs. The forward oligo contains (1) a short (80 bp) arm of homology, (2) the mutation being introduced, and (3) a 3’ sequence which serves as a primer to amplify the large (3 kb) downstream region of homology as depicted. (**D**) Detailed schematic depicting how forward oligos are designed to make deletions (top) and point mutations (bottom).

**Fig. S4.**
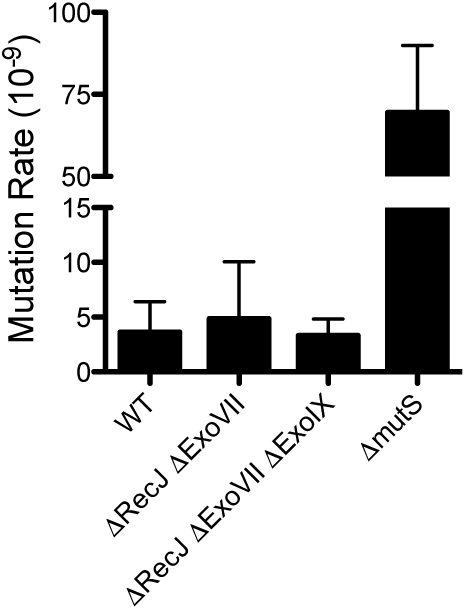
Loss of recJ and exoVII does not increase mutation rate. Fluctuation analysis for spontaneous resistance to rifampicin was performed to assess the mutation rate of the indicated strains. All data are from at least 10 independent biological replicates and shown as the mean±SD.

**Fig. S5.**
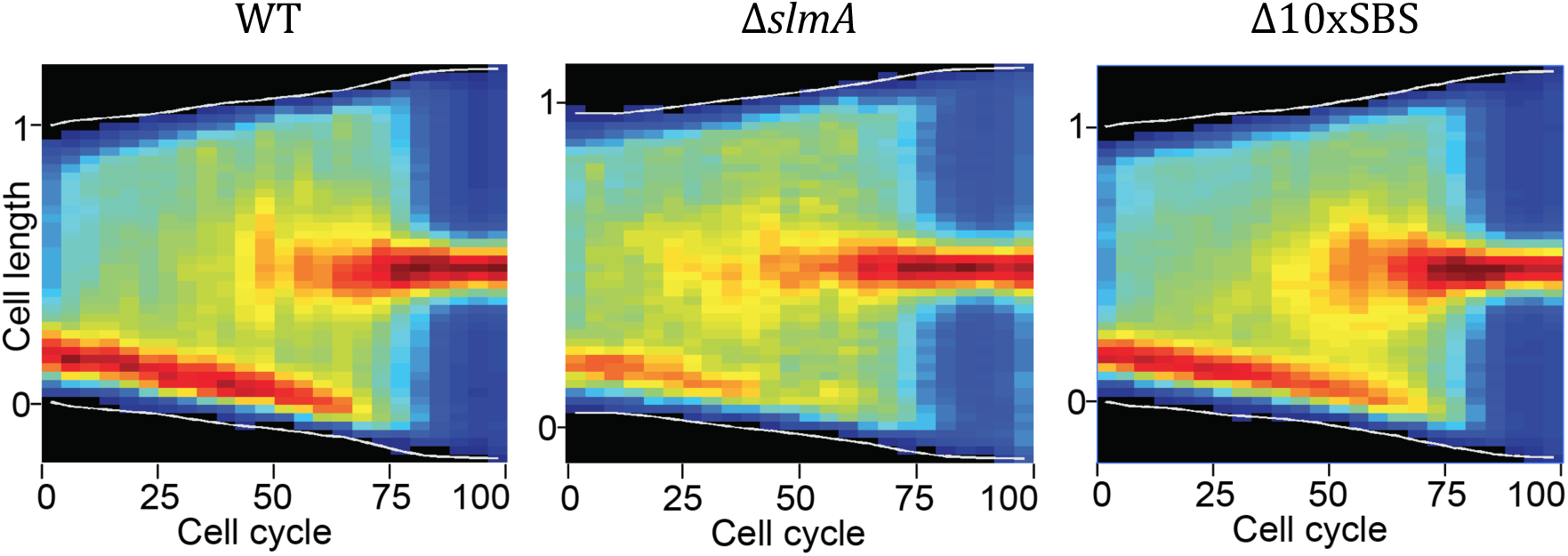
FtsZ localization during the cell cycle is largely unchanged in the Δ10xSBS mutant. Cell cycle choreography for FtsZ-RFPT. Dark red and blue colors were assigned to the maximal and minimal fluorescence intensity projections observed at each time point, respectively. This representation highlights changes in the relative distribution of fluorescence along the long cell axis. Data for each sample are the compilation of data from 40 to 80 single cells.

**Fig. S6.**
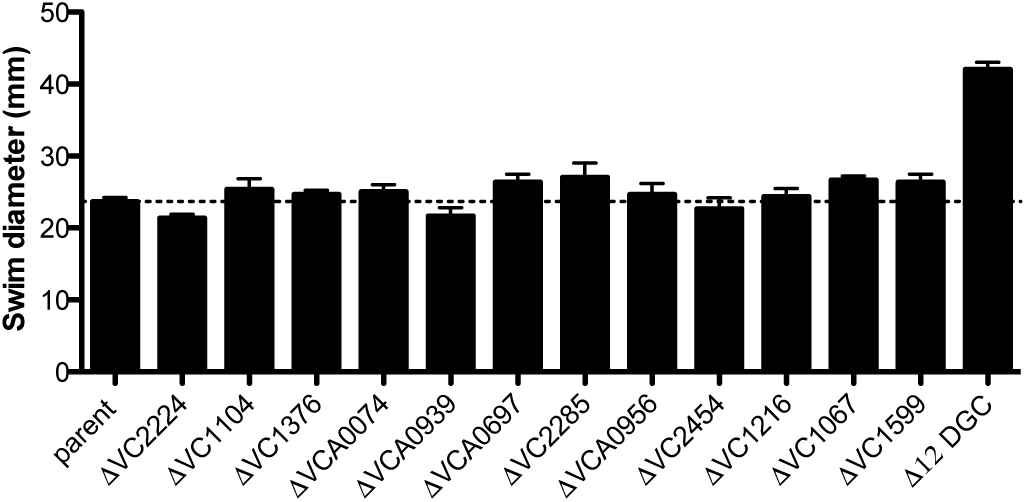
The 12 DGCs targeted act additively or synergistically to reduce swimming motility in V. cholerae. Swim assay performed for the indicated strains. All strains are in a P*_tac_*-*tfoX* Δ*recJ* Δ*exoVII* parent strain background. Data are the result of at least three independent biological replicates and shown as the mean±SD.

## SUPPLEMENTARY TABLES

**Table S1.** RNA-seq expression analysis of all DGCs in *V. cholerae* during growth in rich medium

| Locus<sup>&</sup> | Gene name | Replicate 1<sup>§</sup> | Replicate 2<sup>§</sup> | Replicate 3<sup>§</sup> | Average expression<sup>#</sup> | Expression rank<sup>*</sup> | Biofilm% | Motility% |
| --- | --- | --- | --- | --- | --- | --- | --- | --- |
| VC0072 |  | 0.155 | 0.135 | 0.111 | 0.134 | 20 |  |  |
| VC0130 |  | 0.182 | 0.140 | 0.221 | 0.181 | 16 |  |  |
| VC0398 |  | 0.278 | 0.250 | 0.260 | 0.263 | 10 |  |  |
| VC0653 | rocS | 0.378 | 0.394 | 0.760 | 0.510 | 2 |  | ↓ |
| VC0658 |  | 0.084 | 0.123 | 0.172 | 0.126 | 21 |  |  |
| VC0703 | mbaA | 0.305 | 0.262 | 0.207 | 0.258 | 11 |  |  |
| VC0900 | cdgG | 0.303 | 0.379 | 0.422 | 0.368 | 3 |  |  |
| VC1029 |  | 0.080 | 0.072 | 0.076 | 0.076 | 30 |  |  |
| VC1067 | cdgH | 0.289 | 0.351 | 0.184 | 0.275 | 8 | ↓ | ↑ |
| VC1104 | cdgK | 0.080 | 0.082 | 0.059 | 0.074 | 32 | ↓ | ↑ |
| VC1185 |  | 0.053 | 0.056 | 0.068 | 0.059 | 37 |  |  |
| VC1216 |  | 0.083 | 0.152 | 0.690 | 0.308 | 5 |  |  |
| VC1353 |  | 0.150 | 0.146 | 0.206 | 0.167 | 18 |  |  |
| VC1367 |  | 0.445 | 0.360 | 0.273 | 0.359 | 4 |  |  |
| VC1370 |  | 0.033 | 0.034 | 0.049 | 0.039 | 39 |  |  |
| VC1372 |  | 0.088 | 0.130 | 0.145 | 0.121 | 23 |  |  |
| VC1376 | cdgM | 0.026 | 0.072 | 0.171 | 0.089 | 26 | ↓ |  |
| VC1593 | acgB | 0.048 | 0.029 | 0.119 | 0.065 | 33 |  |  |
| VC1599 |  | 0.216 | 0.246 | 0.340 | 0.267 | 9 |  |  |
| VC1934 |  | 0.030 | 0.023 | 0.066 | 0.040 | 38 |  |  |
| VC2224 |  | 0.087 | 0.070 | 0.073 | 0.077 | 29 |  |  |
| VC2285 | cdgL | 0.318 | 0.235 | 0.309 | 0.287 | 6 | ↓ | ↑ |
| VC2370 |  | 0.032 | 0.041 | 0.105 | 0.059 | 36 |  |  |
| VC2454 | vpvC | 0.141 | 0.180 | 0.248 | 0.189 | 14 | ↓ | ↑ |
| VC2697 |  | 0.016 | 0.004 | 0.042 | 0.020 | 41 |  |  |
| VC2750 |  | 0.054 | 0.041 | 0.139 | 0.078 | 28 |  |  |
| VCA0049 |  | 0.227 | 0.122 | 0.227 | 0.192 | 13 |  |  |
| VCA0074 | cdgA | 0.028 | 0.038 | 0.037 | 0.034 | 40 | ↓ |  |
| VCA0080 |  | 0.081 | 0.121 | 0.168 | 0.123 | 22 |  |  |
| VCA0165 |  | 0.148 | 0.095 | 0.377 | 0.207 | 12 |  |  |
| VCA0217 |  | 0.084 | 0.134 | 0.121 | 0.113 | 24 |  |  |
| VCA0557 |  | 0.152 | 0.085 | 0.269 | 0.169 | 17 |  |  |
| VCA0560 |  | 0.179 | 0.154 | 0.118 | 0.151 | 19 |  |  |
| VCA0697 | cdgD | 0.590 | 0.522 | 0.443 | 0.519 | 1 |  | ↑ |
| VCA0785 | cdgC | 0.041 | 0.029 | 0.110 | 0.060 | 34 |  |  |
| VCA0848 |  | 0.048 | 0.051 | 0.126 | 0.075 | 31 |  |  |
| VCA0939 |  | 0.068 | 0.043 | 0.128 | 0.080 | 27 |  |  |
| VCA0956 |  | 0.159 | 0.171 | 0.220 | 0.183 | 15 |  |  |
| VCA0960 |  | 0.038 | 0.048 | 0.092 | 0.060 | 35 |  |  |
| VCA0965 |  | 0.058 | 0.073 | 0.174 | 0.102 | 25 |  |  |
| VCA1082 |  | 0.219 | 0.296 | 0.319 | 0.278 | 7 |  |  |

<sup>&</sup>The 12 loci highlighted in blue were targeted for inactivation via Exo-MuGENT
<sup>§</sup>Data represent the relative transcript abundance of each gene indicated on the left in three independent biological replicates (see **Methods** for a detailed description on how this analysis was performed)
<sup>#</sup>Average expression is the mean of all three biological replicates.
<sup>\*</sup>Expression of each DGC is indicated on a scale from 1 (highest expression) to 41 (lowest expression)
%Mutants of the genes indicated have previously been observed to display an increase (↑) or decrease (↓) in biofilm formation or motility as indicated (1,2).

**Table S2.**
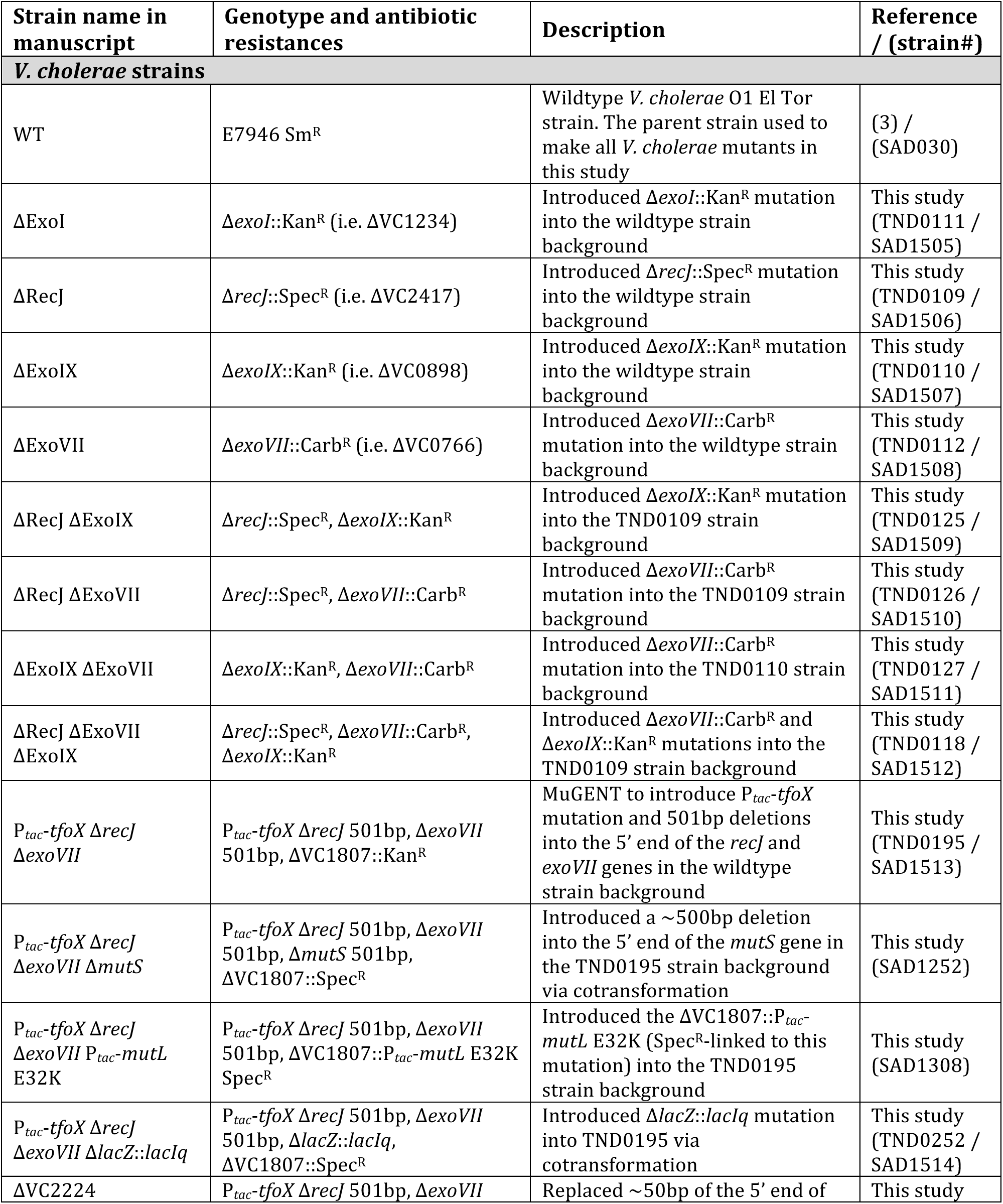
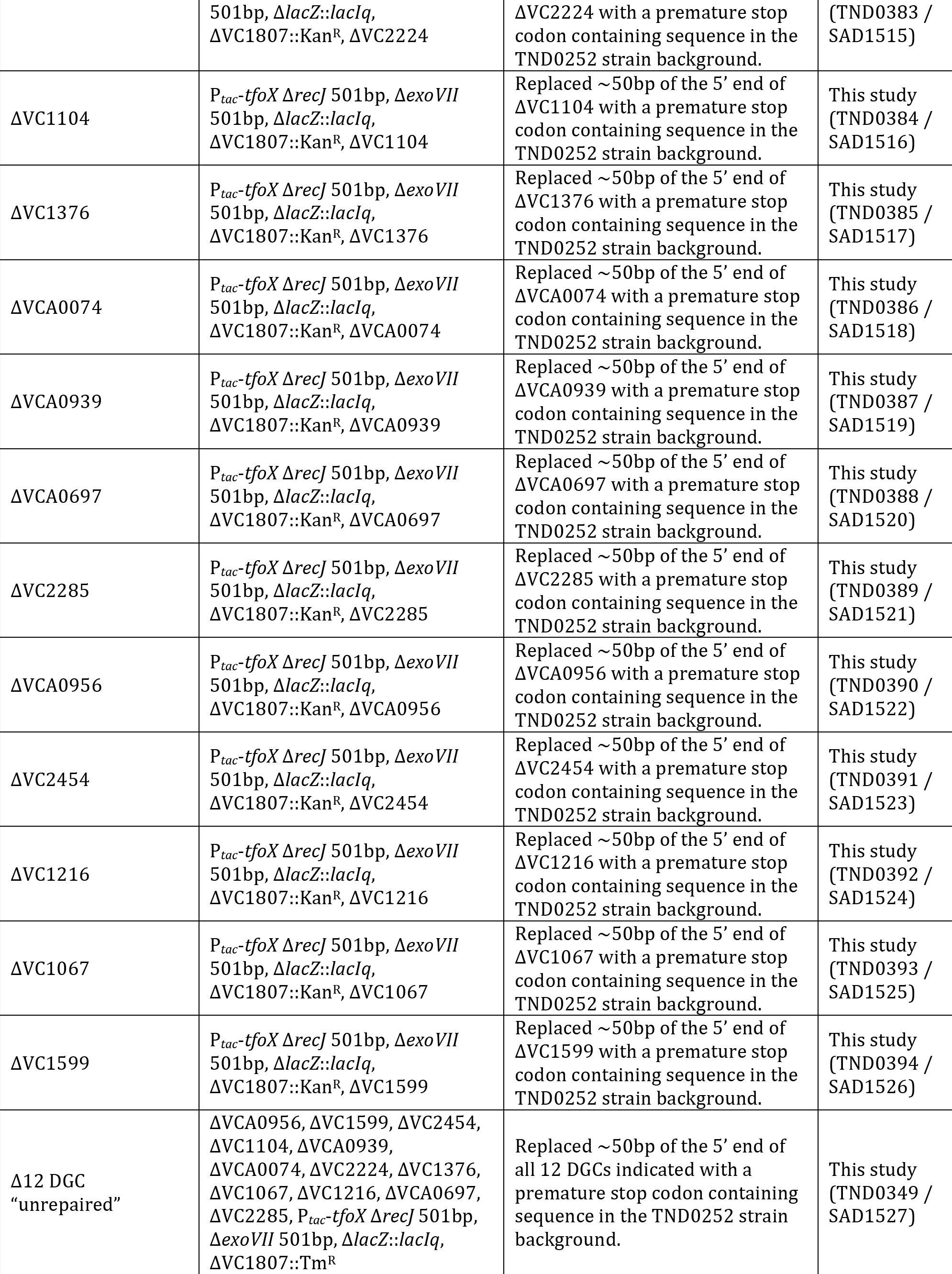
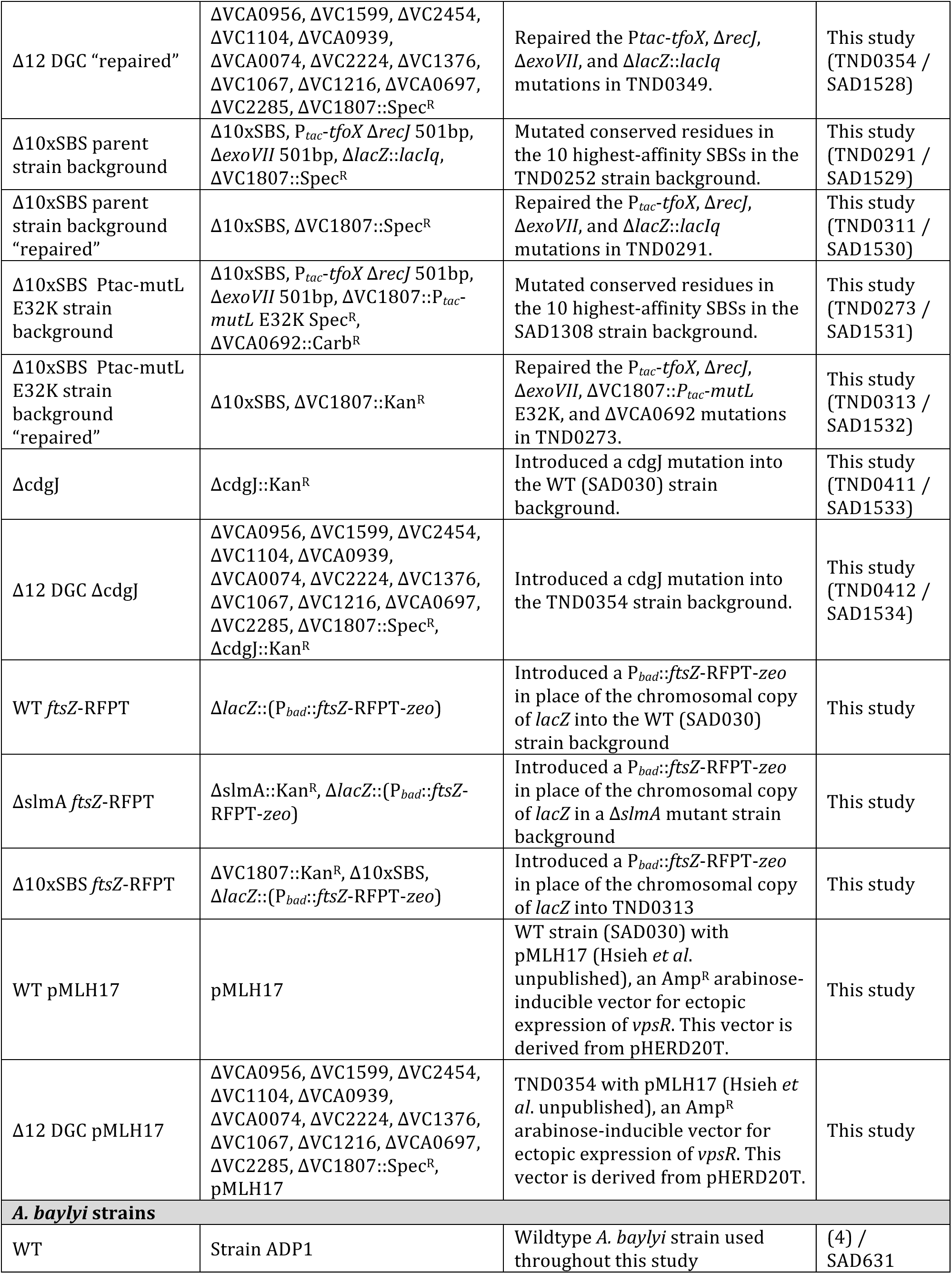
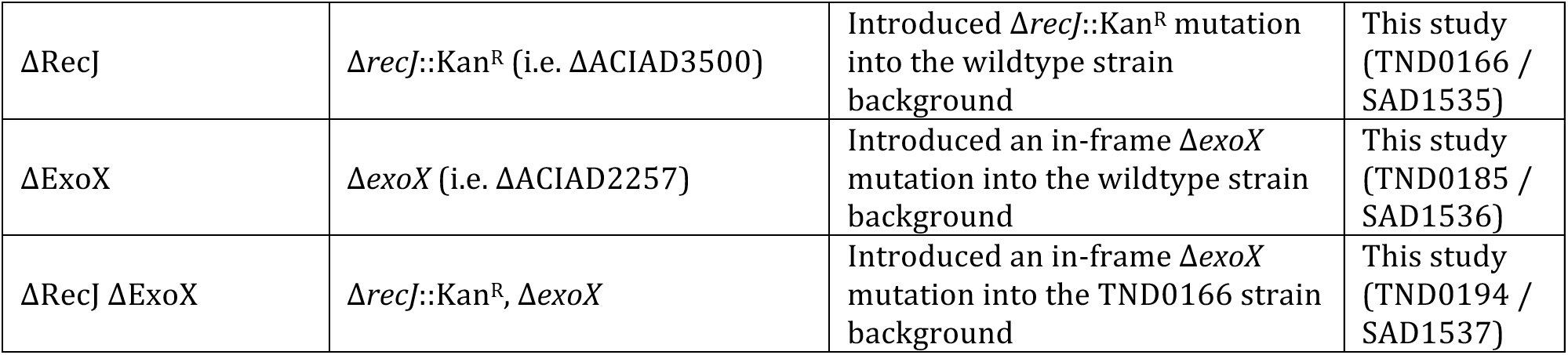
Strains used in this study

**Table S3.**
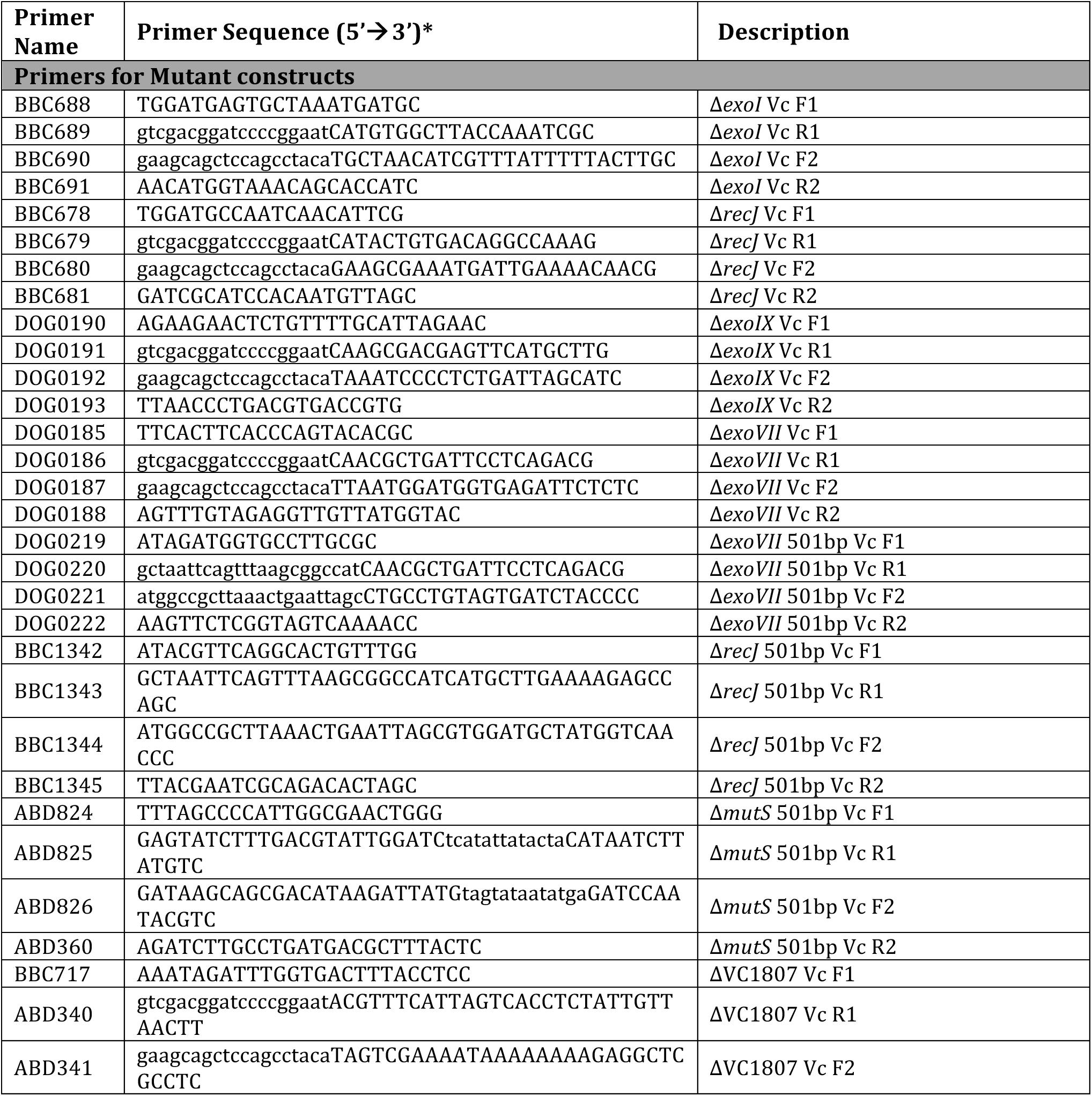
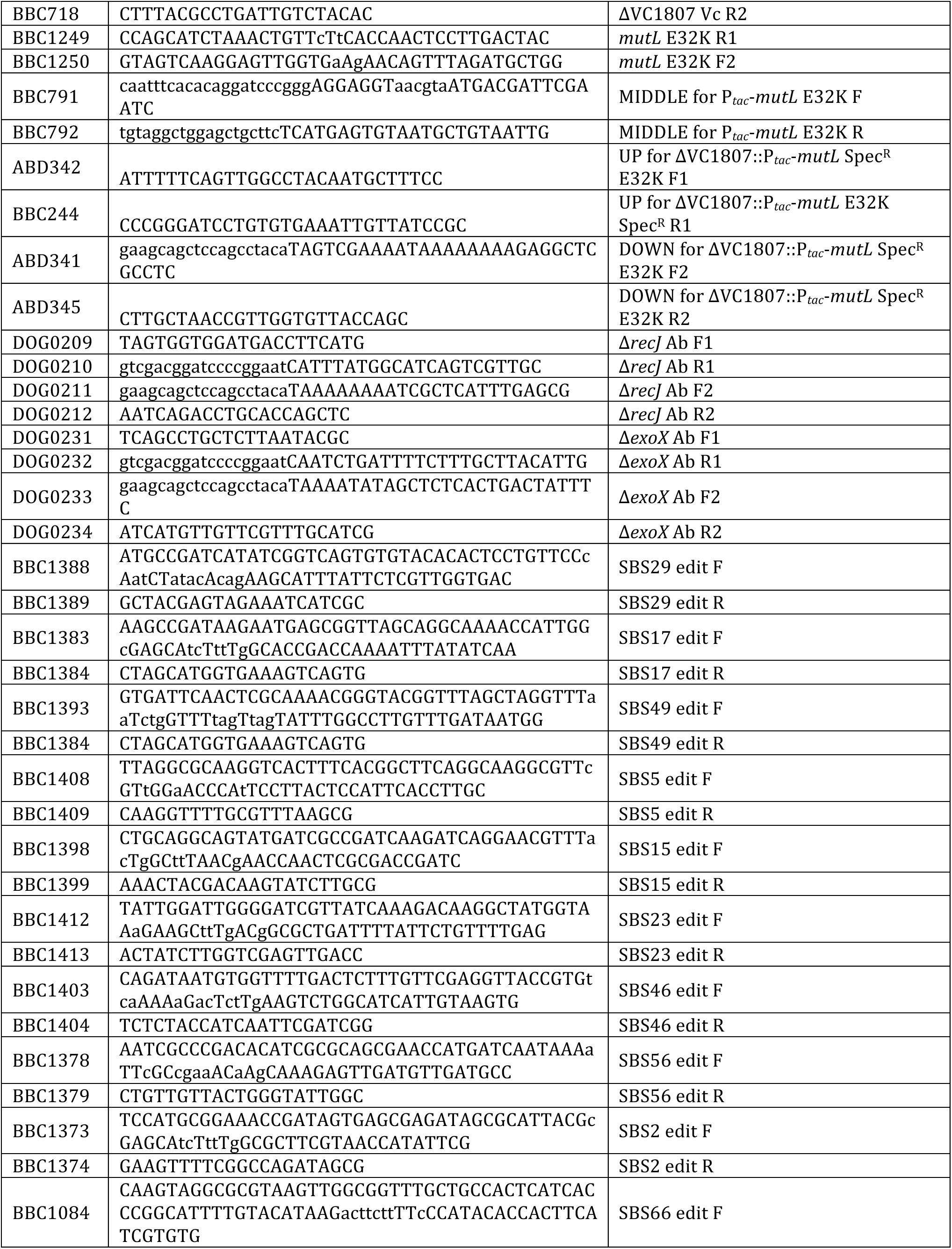
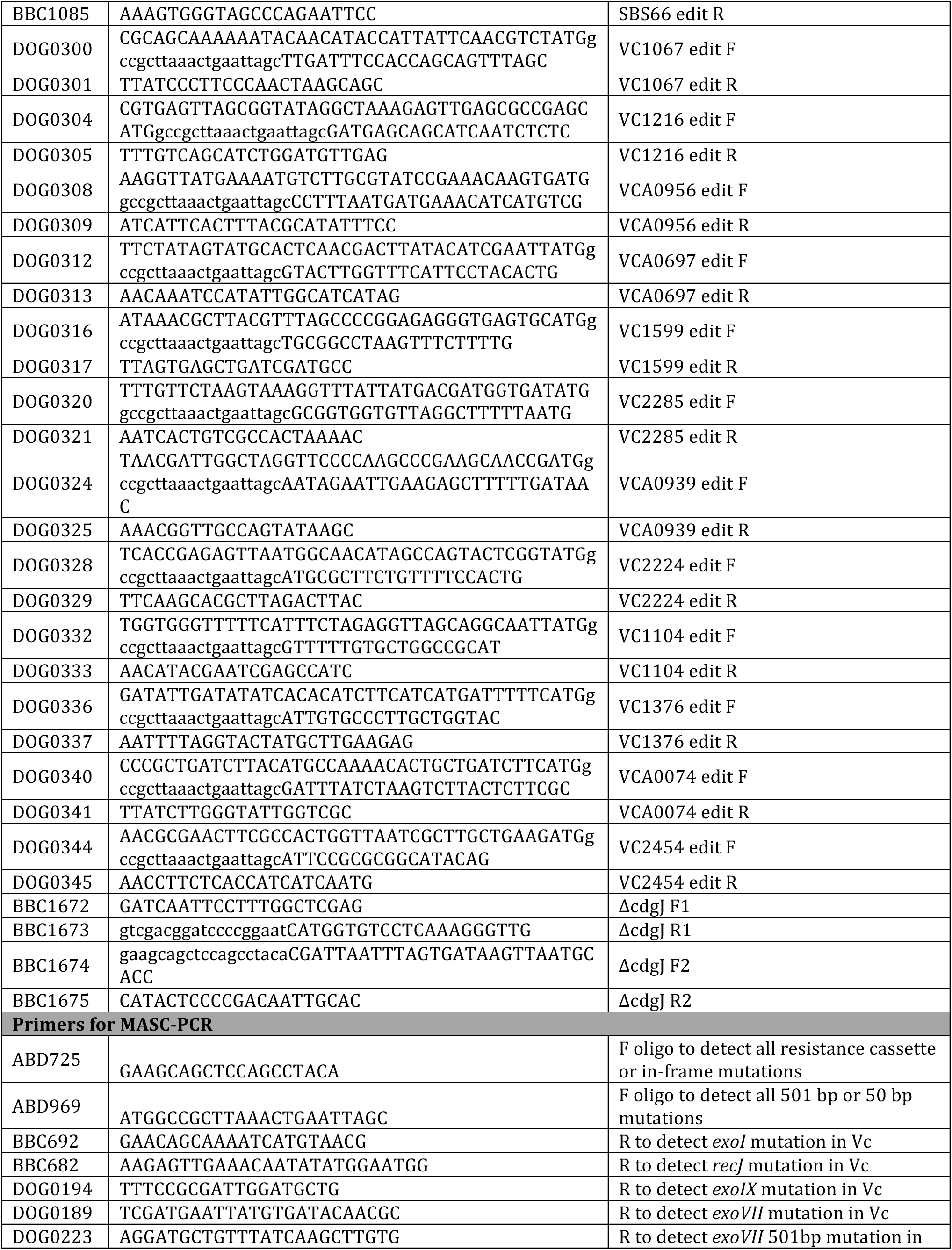
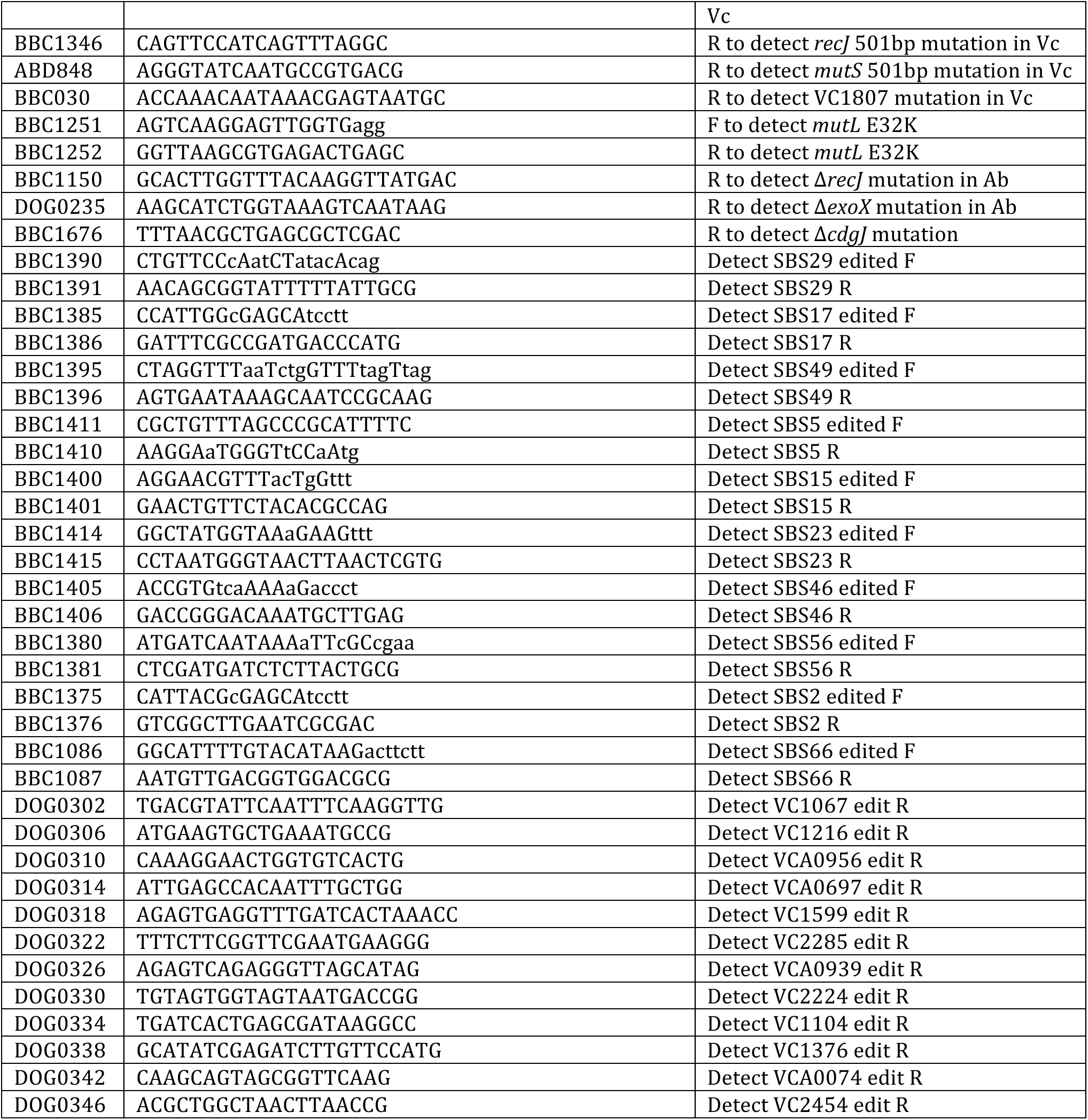
Primers used in this study

## REFERENCES

1. Dalia, A.B., McDonough, E. and Camilli, A. (2014) Multiplex genome editing by natural transformation. Proc Natl Acad Sci U S A, 111, 8937–8942.

2. Watt, V.M., Ingles, C.J., Urdea, M.S. and Rutter, W.J. (1985) Homology requirements for recombination in Escherichia coli. Proc Natl Acad Sci U S A, 82, 4768–4772.

3. Lorenz, M.G. and Wackernagel, W. (1994) Bacterial gene transfer by natural genetic transformation in the environment. Microbiol Rev, 58, 563–602.

4. Miller, V.L., DiRita, V.J. and Mekalanos, J.J. (1989) Identification of toxS, a regulatory gene whose product enhances toxR-mediated activation of the cholera toxin promoter. J Bacteriol, 171, 1288–1293.

5. Juni, E. and Janik, A. (1969) Transformation of Acinetobacter calco-aceticus (Bacterium anitratum). J Bacteriol, 98, 281–288.

6. Blokesch, M. and Schoolnik, G.K. (2008) The extracellular nuclease Dns and its role in natural transformation of Vibrio cholerae. J Bacteriol, 190, 7232–7240.

7. Suckow, G., Seitz, P. and Blokesch, M. (2011) Quorum sensing contributes to natural transformation of Vibrio cholerae in a species-specific manner. J Bacteriol, 193, 4914–4924.

8. Dalia, A.B., Lazinski, D.W. and Camilli, A. (2014) Identification of a membrane-bound transcriptional regulator that links chitin and natural competence in Vibrio cholerae. MBio, 5, e01028–01013.

9. Wang, H.H., Isaacs, F.J., Carr, P.A., Sun, Z.Z., Xu, G., Forest, C.R. and Church, G.M. (2009) Programming cells by multiplex genome engineering and accelerated evolution. Nature, 460, 894–898.

10. Hall, B.M., Ma, C.X., Liang, P. and Singh, K.K. (2009) Fluctuation analysis CalculatOR: a web tool for the determination of mutation rate using Luria-Delbruck fluctuation analysis. Bioinformatics, 25, 1564–1565.

11. Rosche, W.A. and Foster, P.L. (2000) Determining mutation rates in bacterial populations. Methods, 20, 4–17.

12. Lazinski, D.W. and Camilli, A. (2013) Homopolymer tail-mediated ligation PCR: a streamlined and highly efficient method for DNA cloning and library construction. Biotechniques, 54, 25–34.

13. Seed, K.D., Yen, M., Shapiro, B.J., Hilaire, I.J., Charles, R.C., Teng, J.E., Ivers, L.C., Boncy, J., Harris, J.B. and Camilli, A. (2014) Evolutionary consequences of intra-patient phage predation on microbial populations. eLife, 3, e03497.

14. Shishkin, A.A., Giannoukos, G., Kucukural, A., Ciulla, D., Busby, M., Surka, C., Chen, J., Bhattacharyya, R.P., Rudy, R.F., Patel, M.M. et al. (2015) Simultaneous generation of many RNA-seq libraries in a single reaction. Nat Methods, 12, 323–325.

15. Afgan, E., Baker, D., van den Beek, M., Blankenberg, D., Bouvier, D., Cech, M., Chilton, J., Clements, D., Coraor, N., Eberhard, C. et al. (2016) The Galaxy platform for accessible, reproducible and collaborative biomedical analyses: 2016 update. Nucleic Acids Res, 44, W3–W10.

16. Galli, E., Paly, E. and Barre, F.X. (2017) Late assembly of the Vibrio cholerae cell division machinery postpones septation to the last 10% of the cell cycle. Sci Rep, 7, 44505.

17. Galli, E., Poidevin, M., Le Bars, R., Desfontaines, J.M., Muresan, L., Paly, E., Yamaichi, Y. and Barre, F.X. (2016) Cell division licensing in the multi-chromosomal Vibrio cholerae bacterium. Nat Microbiol, 1, 16094.

18. Massie, J.P., Reynolds, E.L., Koestler, B.J., Cong, J.P., Agostoni, M. and Waters, C.M. (2012) Quantification of high-specificity cyclic diguanylate signaling. Proc Natl Acad Sci U S A, 109, 12746–12751.

19. Wolfe, A.J. and Berg, H.C. (1989) Migration of bacteria in semisolid agar. Proc Natl Acad Sci U S A, 86, 6973–6977.

20. Sambanthamoorthy, K., Gokhale, A.A., Lao, W., Parashar, V., Neiditch, M.B., Semmelhack, M.F., Lee, I. and Waters, C.M. (2011) Identification of a novel benzimidazole that inhibits bacterial biofilm formation in a broad-spectrum manner. Antimicrob Agents Chemother, 55, 4369–4378.

21. Marvig, R.L. and Blokesch, M. (2010) Natural transformation of Vibrio cholerae as a tool–optimizing the procedure. BMC Microbiol, 10, 155.

22. Burdett, V., Baitinger, C., Viswanathan, M., Lovett, S.T. and Modrich, P. (2001) In vivo requirement for RecJ, ExoVII, ExoI, and ExoX in methyl-directed mismatch repair. Proc Natl Acad Sci U S A, 98, 6765–6770.

23. Claverys, J.P. and Lacks, S.A. (1986) Heteroduplex deoxyribonucleic acid base mismatch repair in bacteria. Microbiol Rev, 50, 133–165.

24. Claverys, J.P., Mejean, V., Gasc, A.M. and Sicard, A.M. (1983) Mismatch repair in Streptococcus pneumoniae: relationship between base mismatches and transformation efficiencies. Proc Natl Acad Sci U S A, 80, 5956–5960.

25. Nyerges, A., Csorgo, B., Nagy, I., Latinovics, D., Szamecz, B., Posfai, G. and Pal, C. (2014) Conditional DNA repair mutants enable highly precise genome engineering. Nucleic Acids Res.

26. Humbert, O., Prudhomme, M., Hakenbeck, R., Dowson, C.G. and Claverys, J.P. (1995) Homeologous recombination and mismatch repair during transformation in Streptococcus pneumoniae: saturation of the Hex mismatch repair system. Proc Natl Acad Sci U S A, 92, 9052–9056.

27. Hayes, C.A., Dalia, T.N. and Dalia, A.B. (2017) Systematic genetic dissection of PTS in Vibrio cholerae uncovers a novel glucose transporter and a limited role for PTS during infection of a mammalian host. Mol Microbiol.

28. Romling, U., Galperin, M.Y. and Gomelsky, M. (2013) Cyclic di-GMP: the first 25 years of a universal bacterial second messenger. Microbiol Mol Biol Rev, 77, 1–52.

29. D'Souza, M., Glass, E.M., Syed, M.H., Zhang, Y., Rodriguez, A., Maltsev, N. and Galperin, M.Y. (2007) Sentra: a database of signal transduction proteins for comparative genome analysis. Nucleic Acids Res, 35, D271–273.

30. Liu, X., Beyhan, S., Lim, B., Linington, R.G. and Yildiz, F.H. (2010) Identification and characterization of a phosphodiesterase that inversely regulates motility and biofilm formation in Vibrio cholerae. J Bacteriol, 192, 4541–4552.

31. Townsley, L. and Yildiz, F.H. (2015) Temperature affects c-di-GMP signalling and biofilm formation in Vibrio cholerae. Environ Microbiol, 17, 4290–4305.

32. Waters, C.M., Lu, W., Rabinowitz, J.D. and Bassler, B.L. (2008) Quorum sensing controls biofilm formation in Vibrio cholerae through modulation of cyclic di-GMP levels and repression of vpsT. J Bacteriol, 190, 2527–2536.

33. Hengge, R. (2009) Principles of c-di-GMP signalling in bacteria. Nat Rev Microbiol, 7, 263–273.

34. Yildiz, F.H., Dolganov, N.A. and Schoolnik, G.K. (2001) VpsR, a Member of the Response Regulators of the Two-Component Regulatory Systems, Is Required for Expression of vps Biosynthesis Genes and EPS(ETr)-Associated Phenotypes in Vibrio cholerae O1 El Tor. J Bacteriol, 183, 1716-1726.

35. Srivastava, D., Harris, R.C. and Waters, C.M. (2011) Integration of cyclic di-GMP and quorum sensing in the control of vpsT and aphA in Vibrio cholerae. J Bacteriol, 193, 6331–6341.

36. Krasteva, P.V., Fong, J.C., Shikuma, N.J., Beyhan, S., Navarro, M.V., Yildiz, F.H. and Sondermann, H. (2010) Vibrio cholerae VpsT regulates matrix production and motility by directly sensing cyclic di-GMP. Science, 327, 866–868.

37. Jones, C.J., Utada, A., Davis, K.R., Thongsomboon, W., Zamorano Sanchez, D., Banakar, V., Cegelski, L., Wong, G.C. and Yildiz, F.H. (2015) C-di-GMP Regulates Motile to Sessile Transition by Modulating MshA Pili Biogenesis and Near-Surface Motility Behavior in Vibrio cholerae. PLoS Pathog, 11, e1005068.

38. Harms, K., Schon, V., Kickstein, E. and Wackernagel, W. (2007) The RecJ DNase strongly suppresses genomic integration of short but not long foreign DNA fragments by homology-facilitated illegitimate recombination during transformation of Acinetobacter baylyi. Mol Microbiol, 64, 691–702.

39. Mosberg, J.A., Gregg, C.J., Lajoie, M.J., Wang, H.H. and Church, G.M. (2012) Improving lambda red genome engineering in Escherichia coli via rational removal of endogenous nucleases. PLoS One, 7, e44638.

40. Chase, J.W. and Richardson, C.C. (1977) Escherichia coli mutants deficient in exonuclease VII. J Bacteriol, 129, 934–947.

41. Viswanathan, M. and Lovett, S.T. (1998) Single-strand DNA-specific exonucleases in Escherichia coli. Roles in repair and mutation avoidance. Genetics, 149, 7–16.

42. Dalia, T.N., Hayes, C.A., Stoylar, S., Marx, C.J., Mckinlay, J.B. and Dalia, A.B. (2017) Multiplex genome editing by natural transformation (MuGENT) for synthetic biology in Vibrio natriegens. bioRxiv, doi:10.1101/122655.

43. Bubendorfer, S., Krebes, J., Yang, I., Hage, E., Schulz, T.F., Bahlawane, C., Didelot, X. and Suerbaum, S. (2016) Genome-wide analysis of chromosomal import patterns after natural transformation of Helicobacter pylori. Nat Commun, 7, 11995.

## SUPPLEMENTARY REFERENCES

1. Liu, X., Beyhan, S., Lim, B., Linington, R.G. and Yildiz, F.H. (2010) Identification and characterization of a phosphodiesterase that inversely regulates motility and biofilm formation in Vibrio cholerae. J Bacteriol, 192, 4541–4552.

2. Townsley, L. and Yildiz, F.H. (2015) Temperature affects c-di-GMP signalling and biofilm formation in Vibrio cholerae. Environ Microbiol, 17, 4290–4305.

3. Miller, V.L., DiRita, V.J. and Mekalanos, J.J. (1989) Identification of toxS, a regulatory gene whose product enhances toxR-mediated activation of the cholera toxin promoter. J Bacteriol, 171, 1288–1293.

4. Juni, E. and Janik, A. (1969) Transformation of Acinetobacter calco-aceticus (Bacterium anitratum). J Bacteriol, 98, 281–288.

